# Regulation of mitophagy by the NSL complex underlies genetic risk for Parkinson’s disease at Chr16q11.2 and on the MAPT H1 allele

**DOI:** 10.1101/2020.01.06.896241

**Authors:** Marc P.M. Soutar, Daniela Melandri, Benjamin O’Callaghan, Emily Annuario, Amy E. Monaghan, Natalie J. Welsh, Karishma D’Sa, Sebastian Guelfi, David Zhang, Alan Pittman, Daniah Trabzuni, Kylie S. Pan, Demis A. Kia, Magda Bictash, Sonia Gandhi, Henry Houlden, Mark R. Cookson, Nicholas W Wood, Andrew B. Singleton, John Hardy, Paul J. Whiting, Cornelis Blauwendraat, Alexander J. Whitworth, Claudia Manzoni, Mina Ryten, Patrick A. Lewis, Hélène Plun-Favreau

**Affiliations:** UCL Queen Square Institute of Neurology, London, UK; King’s College, London, UK; UCL Alzheimer’s Research UK, Drug Discovery Institute, London, UK; UCL Dementia Research Institute, London, UK; MRC Mitochondrial Biology Unit, University of Cambridge, Cambridge, UK; Francis Crick Institute, London, UK; UCL NIHR Great Ormond Street Hospital, London, UK; St Georges University, London, UK; Laboratory of Neurogenetics, National Institute on Aging, National Institutes of Health, Bethesda, MD, USA; School of Pharmacy, University of Reading, Reading, UK; Royal Veterinary College, London, UK

**Author notes:** these authors contributed equally to the work (Marc P.M. Soutar, Daniela Melandri, Benjamin O’Callaghan, Emily Annuario). these authors contributed equally to the work (Mina Ryten, Patrick A. Lewis, Hélène Plun-Favreau).

**Keywords:** GWAS, KANSL1, KAT8, mitophagy, Parkinson’s disease

## Abstract

Parkinson’s disease (PD) is a common incurable neurodegenerative disease. The identification of genetic variants via genome-wide association studies (GWAS) has considerably advanced our understanding of the PD genetic risk. Understanding the functional significance of the risk loci is now a critical step towards translating these genetic advances into an enhanced biological understanding of the disease. Impaired mitophagy is a key causative pathway in familial PD, but its relevance to idiopathic PD is unclear. We used a mitophagy screening assay to evaluate the functional significance of risk genes identified through GWAS. We identified two new regulators of PINK1-mitophagy, KAT8 and KANSL1, previously shown to modulate lysine acetylation. We show that KAT8 and KANSL1 modulate *PINK1* gene expression and subsequent PINK1-mitophagy. These findings suggest PINK1-mitophagy is a contributing factor to idiopathic PD. *KANSL1* is located on chromosome 17q21 where the risk associated gene has long been considered to be *MAPT*. While our data does not exclude a possible association between the *MAPT* gene and PD, it provides strong evidence that *KANSL1* plays a crucial role in the disease. Finally, these results enrich our understanding of physiological events regulating mitophagy and establish a novel pathway for drug targeting in neurodegeneration.

## INTRODUCTION

Parkinson’s disease (PD) is the most common movement disorder of old age and afflicts more than 125,000 in the UK (Hardy *et al*., 2009). Temporary symptomatic relief remains the cornerstone of current treatments, with no disease-modifying therapies yet available (Connolly and Lang, 2014). Until recently, the genetic basis for PD was limited to family-based linkage studies, favouring the identification of rare Mendelian genes of high penetrance and effect. However, genome-wide association studies (GWAS) have identified large numbers of common genetic variants linked to increased risk of developing the disease (Chang *et al*., 2017; Nalls *et al*., 2019). While these genetic discoveries have led to a rapid improvement in our understanding of the genetic architecture of PD (Nalls *et al*., 2011), they have resulted in two major challenges for the research community. First, conclusively identifying the causal gene(s) for a given risk locus, and secondly dissecting their contribution to disease pathogenesis. Addressing these challenges is critical for moving beyond genetic insights to developing new disease-modifying strategies for PD.

Previous functional analyses of *PINK1* and *PRKN*, two genes associated with autosomal recessive PD, have highlighted the selective degradation of damaged mitochondria (mitophagy) as a key contributor to disease pathogenesis. In mammalian cells, the mitochondrial kinase PINK1 selectively accumulates at the surface of damaged mitochondria, where it phosphorylates ubiquitin, leading to the recruitment and phosphorylation of the E3 ubiquitin ligase Parkin. The recruitment of autophagy receptors leads to the engulfment of damaged mitochondria in autophagosomes, and ultimately fusion with lysosomes (Narendra *et al*., 2008, 2010; Kazlauskaite *et al*., 2014; Shiba-Fukushima *et al*., 2014; Lazarou *et al*., 2015; McWilliams and Muqit, 2017). It has subsequently become clear that other PD-associated Mendelian genes, such as *FBXO7, DJ-1* and *VPS35* (Plotegher and Duchen, 2017), are implicated in the regulation of PINK1-mediated mitochondrial quality control. Based upon these data, we hypothesised that PD-GWAS candidate genes may also be involved in this process, providing a mechanistic link between these genes and the aetiology of idiopathic PD. In order to test that hypothesis, we used functional genomics to prioritise candidate genes at the PD GWAS loci, and we developed a biological screening assay as a tool to identify genes that regulate PINK1-mitophagy, and as such, are very likely to be genes that increase the risk of developing PD.

In this study, we show that *KAT8* and *KANSL1*, two genes that were previously shown to be part of the same lysine acetylase complex partially located at the mitochondria (Chatterjee *et al*., 2016), are new and important regulators of *PINK1* gene transcription and PINK1-mediated mitochondrial quality control. These findings suggest mitophagy contributes to idiopathic PD and provides a proof of principle for functional screening approaches to identify causative genes in GWAS loci. Finally, these results suggest lysine acetylation as a potential new avenue for mitophagy modulation and therapeutic intervention.

## RESULTS

Genomic analyses of PD have identified over 80 loci associated with an increased lifetime risk for disease (Chang *et al*., 2017). In contrast to Mendelian PD genes, however, the assignment of a causative gene to a risk locus is often challenging. In order to identify new risk genes for PD, we undertook a triage of PD GWAS candidate genes using a combination of methods: i) Colocalization (Coloc) and transcriptome-wide association analysis (TWAS) (Giambartolomei *et al*., 2014) using expression quantitative trait loci (eQTLs) information derived from Braineac (Ramasamy *et al*., 2014), GTEx and CommonMind resources (Lonsdale *et al*., 2013; Kia *et al*., 2019) to link PD risk variants with specific genes, ii) weighted protein-protein interaction (PPI) network analysis (WPPINA)(Ferrari *et al*., 2018) based on Mendelian genes associated with PD, and iii) the prioritised gene set as described in PD-GWAS (Nalls *et al*., 2014; Chang *et al*., 2017). This resulted in the nomination of 31 open reading frames (ORFs) as putatively causal for associations at PD risk loci. 55% of these genes were prioritised through multiple techniques, with three out of 31 genes (*KAT8, CTSB* and *NCKIPSD*) identified through all three prioritization methods (Extended Data Fig. 1A). The 31 genes, together with 7 PD Mendelian genes and lysosomal storage disorder genes, previously shown to be enriched for rare, likely damaging variants in PD (Robak *et al*., 2017), were then taken forward for functional analysis (Fig. 1A).

**Figure 1.**
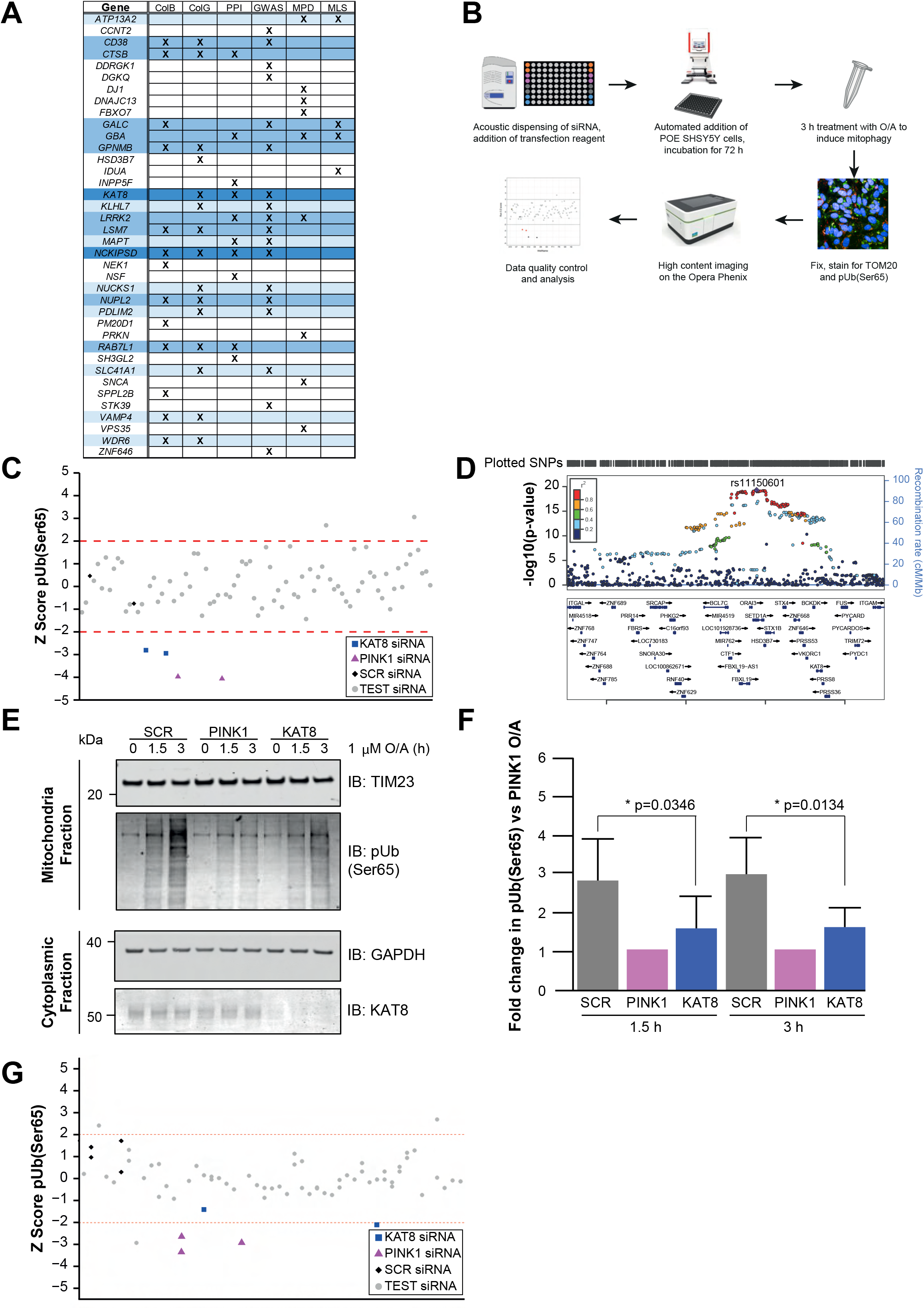
High content mitophagy screen of PD risk genes identifies KAT8 as a modulator of pUb(Ser65) levels. **A**. The heat-map represents increasing evidence for gene prioritization (white, light blue, and dark blue: one, two, and three evidences, respectively). ColB = coloc analysis using Braineac, ColG = coloc analysis using GTEx, WPPINA = weighted protein interaction network; GWAS = genes prioritised in PD-GWAS (Chang *et al*., 2017), MPD = Mendelian genes associated with PD, MLS = Mendelian genes associated with lysosomal storage disorders. **B**. Workflow of the high content screen for O/A-induced pUb(Ser65) levels. **C**. pUb(Ser65) Z-scores of one representative mitophagy screen plate. **D**. Overview of the PD GWAS genetic signal at the *KAT8* locus. **E**. Representative IB of mitochondrial fractions from SCR, PINK1 and KAT8 KD POE SH-SY5Y treated with 1 µM O/A for 1.5 or 3 h. **F**.Quantification of pUb(Ser65) in E (n=5, one-way ANOVA with Dunnett’s correction). **G**. pUb(Ser65) Z-scores of one representative KAT screen plate. See Supplementary Table 3 for the complete list of the genes screened. Data are shown as mean ± SD.

Based upon extensive data implicating impaired mitophagy in the aetiology of familial PD, we hypothesized that additional PD-GWAS candidate genes, involved in the most common, idiopathic form of the disease, may play a role in this process. In order to test whether the 38 prioritised genes have a role in PINK1-mitophagy, we developed and optimized a high content screening (HCS) assay for phosphorylation of ubiquitin at serine 65 (pUb(Ser65)), a PINK1-dependent mitophagy marker (Hou *et al*., 2018), following mitochondrial depolarization (Fig. 1B). The 38 prioritised genes were individually knocked down (KD) using siRNA in Parkin over-expressing (POE)-SHSY5Y human neuroblastoma cells. Increased mitochondrial clearance following mitochondrial depolarization induced by treatment with 10 µM of oligomycin/antimycin A (O/A) was validated as an endpoint for mitophagy (Extended Data Fig. 1B). Over 97% of the pUb(Ser65) signal colocalised with the TOM20 mitochondrial marker in O/A treated cells (Extended Data Fig. 1C, D). siRNA KD efficiency was validated using both a pool of *PINK1* siRNA, which decreased O/A induced pUb(Ser65) and subsequent TOM20 degradation (Extended Data Fig. 1E-G) without decreasing cell viability (Extended Data Fig. 2A-B), and a pool of Polo-like kinase 1 (PLK-1) siRNA that decreased cell viability by apoptosis (Extended Data Fig. 2A-B). The siRNA pools for the 38 candidate genes, together with controls, were screened in duplicate on each plate, across three replicate plates per run. Hits were identified as those wells where O/A-induced pUb(Ser65) was decreased or increased at greater than two standard deviations from the mean of the scramble (SCR) negative control siRNA.

*KAT8* was selected based on reproducible downregulation of O/A-induced PINK1-dependent pUb(Ser65) across all three replicates (Fig. 1C and Extended Data Fig. 1H), without affecting cell viability (Extended Data Fig. 2C). Notably, *KAT8* was selected as a candidate gene on the basis of all three prioritization criteria – namely, proximity of the lead SNP to an ORF (Fig 1D), colocalization of a brain-derived eQTL signal with a PD GWAS association signal (Extended Data Fig. 3) and evidence of PPI with a known PD gene (Fig. 1A). Furthermore, we find that colocalization and TWAS (Gusev *et al*., 2016) analyses at this locus are consistent with the KD models in the HCS assay (Supplementary Tables 1 and 2)(Kia *et al*., 2019). Both methods predict that the risk allele operates by reducing *KAT8* expression in PD cases versus controls. The effect of KAT8 KD on pUb(Ser65) was further validated in POE SHSY5Y cells treated with 1 µM O/A, using both immunoblotting (IB) and immunofluorescence (IF) (Fig. 1E-F and Extended Data Fig. 4). In order to assess whether other lysine acetyltransferases (KATs) could regulate PINK1-dependent mitophagy, the pUb(Ser65) screen was repeated in POE SHSY5Y cells silenced for 22 other KATs (Simon *et al*., 2016; Sheikh and Akhtar, 2019). Only *KAT8* KD led to a decreased pUb(Ser65) signal, emphasising the specificity of the KAT8 KD effect on pUb(Ser65) (Fig. 1G and Supplementary Table 3).

These functional data complement and support the omic prioritization of *KAT8* as a causative gene candidate for the chromosome 16q11.2 PD associated locus (Fig. 1D). To gain further insight into a possible role for KAT8 in the aetiology of PD, we explored the known functional interactions of this protein. KAT8 has previously been shown to partially localise to mitochondria as part of the NSL complex together with KANSL1, KANSL2, KANSL3, and MCRS1 (Chatterjee *et al*., 2016). To test whether other components of the NSL complex also modulate mitophagy, the pUb(Ser65) screen was repeated in POE SHSY5Y cells silenced for each of the nine NSL components (HCFC1, KANSL1, KANSL2, KANSL3, KAT8, MCRS1, OGT, PHF20, WDR5). Notably, reduction of KANSL1, KANSL2, KANSL3, MCRS1 and KAT8 expression led to decreased pUb(Ser65) after 1.5 or 3 h O/A treatment (Fig. 2A and Extended Data Fig. 5). Interestingly, *KANSL1* is another PD GWAS candidate gene (Chang *et al*., 2017). The effect of KANSL1 KD on pUb(Ser65) was further validated in POE SHSY5Y cells treated with 1 µM O/A, using both IF and IB (Fig. 2B-E). The effect of the KAT8 and KANSL1 KD on pUb(Ser65) was confirmed in WT SHSY5Y cells expressing endogenous levels of Parkin, and in the astroglioma H4 cell line (Extended Data Fig. 6). In order to further assess the effect of KAT8 and KANSL1 KD on PINK1 activity, we measured pUb(Ser65) levels over time (Fig. 3A-B), as well as Parkin recruitment (Fig. 3C-D) and phosphorylation at Ser65 (pParkin(Ser65)) (Fig. 3E-F) (Kazlauskaite *et al*., 2014). While individual KD of either KANSL1 or KAT8 affect phosphorylation (mean % of pParkin(Ser65) positive mitochondria in O/A-treated cells: SCR 19.601±3.927, PINK1 10.426 ±4.083, KANSL1 12.929±3.214, KAT8 17.115±3.688) and recruitment (mean ratio of mitochondrial FLAG intensity in O/A treated cells: SCR 2.185±0.232, PINK1 1.485±0.222, KANSL1 1.672±0.187, KAT8 2.069±0.213) of Parkin, KANSL1 KD decreased PINK1-dependent activity more efficiently than KAT8 KD (Fig. 3). KD of both KAT8 and KANSL1 reduced subsequent mitochondrial clearance in live POE-SHSY5Y cells, as measured by the mitophagy reporter mt-Keima (Katayama *et al*., 2011) (Fig 4). In order to assess the role of KAT8/KANSL1 in neuronal function and survival *in vivo*, we used *Drosophila* as a simple model system. Notably, the NSL complex was originally discovered in *Drosophila* through the homologs of *KAT8* and *KANSL1* (*mof* and *nsl1*, respectively), but null mutations for these genes are associated with developmental lethality owing to profound transcriptional remodelling during development (Raja *et al*., 2010). Therefore, we utilised inducible transgenic RNAi strains to target the KD of *mof* and *nsl1* specifically in neuronal tissues. Using behavioural assays as a sensitive readout of neuronal function we found that pan-neuronal KD of *mof* or *nsl1* caused progressive loss of motor (climbing) ability (Extended Data Fig. 7A, B), and also significantly shortened lifespan (Extended Data Fig. 7C, D). Interestingly, loss of *nsl1* had a notably stronger effect than loss of *mof*. Consistent with this, KD of *nsl1* but not *mof*, in either all neurons or only in dopaminergic (DA) neurons, caused the loss of DA neurons (Extended Data Fig. 7E, F).

**Figure 2.**
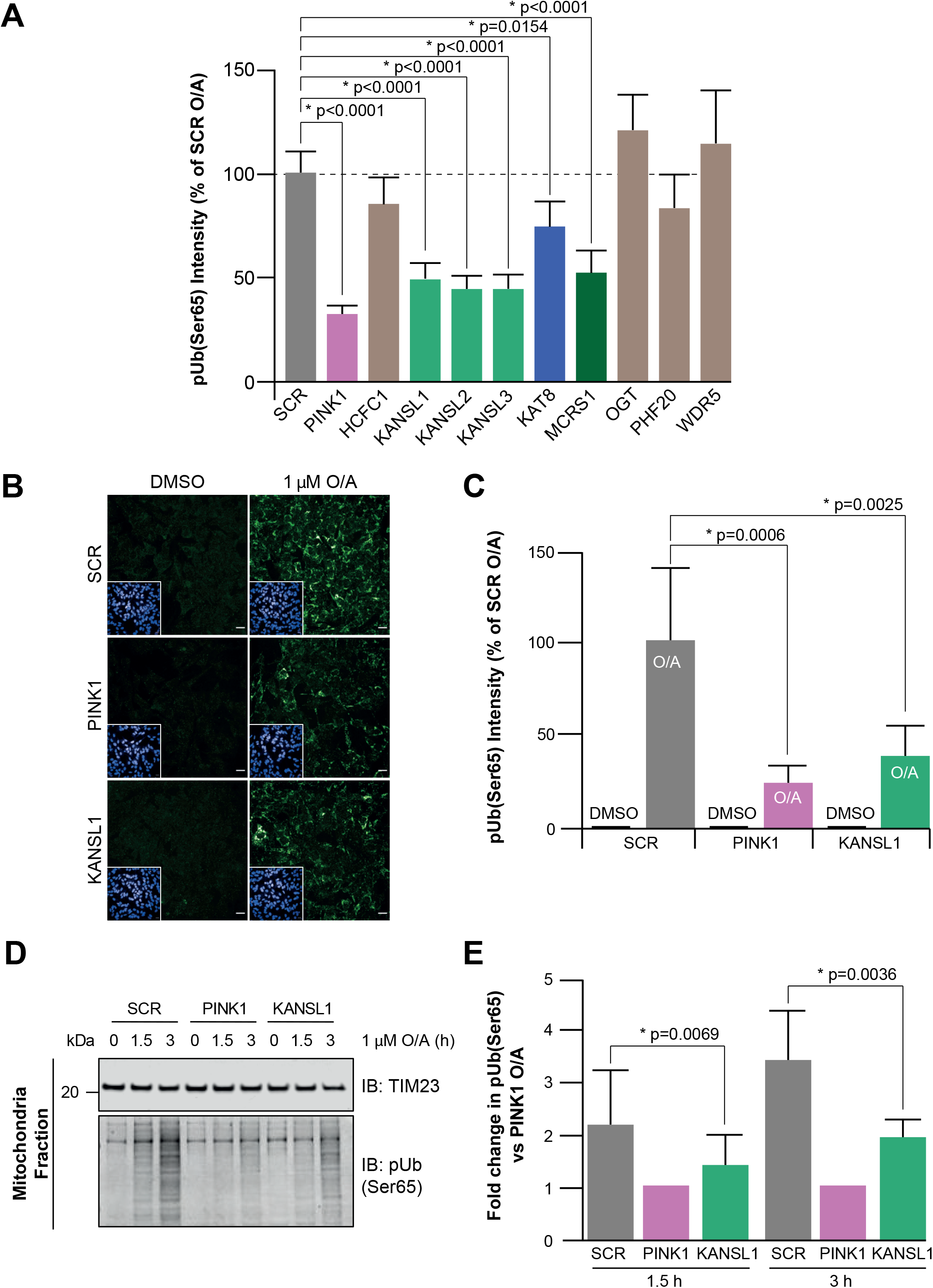
KANSL1, another PD risk gene, also affects pUb(Ser65) levels. **A**. Quantification of pUb(Ser65) following treatment of SCR, PINK1 or NSL components siRNA KD POE SH-SY5Y cells with 1 μM O/A for 1.5 h. Data are shown as mean ± SD; n=6, one-way ANOVA with Dunnett’s correction. **B**. Representative images of pUb(Ser65) following treatment of SCR, PINK1 and KANSL1 KD POE SH-SY5Y cells with 1 µM O/A for 3 h. Insets show the nuclei for the same fields. Scale bar: 20 μm. **C**.Quantification of pUb(Ser65) in B (n=3, two-way ANOVA with Dunnett’s correction). **D**. Representative IB of mitochondrial fractions from SCR, PINK1 and KANSL1 KD POE SH-SY5Y treated with 1 μM O/A for 1.5 or 3 h. **E**. Quantification of pUb(Ser65) in D (n=5, one-way ANOVA with Dunnett’s correction). Data are shown as mean ± SD.

**Figure 3.**
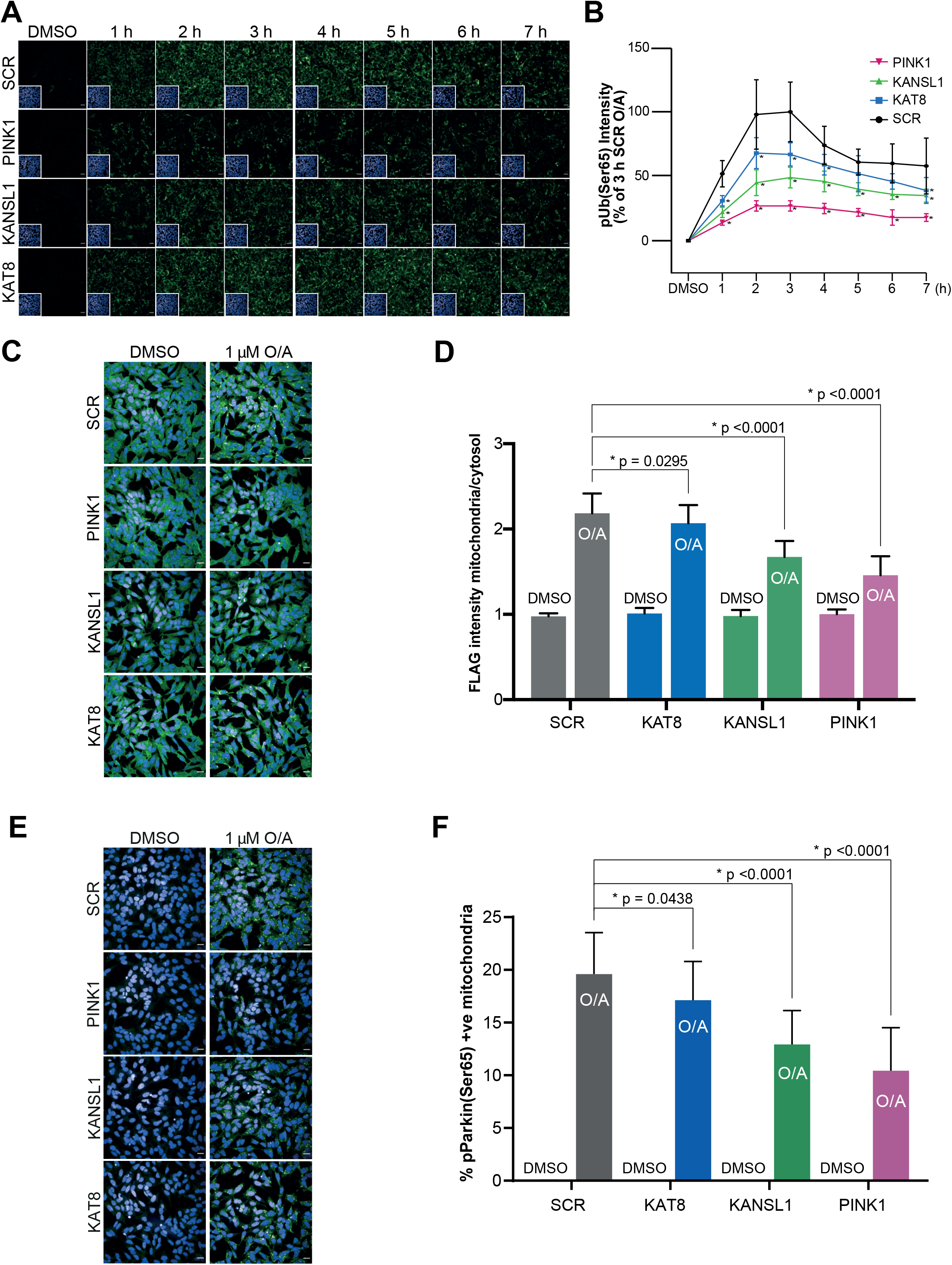
KAT8 and KANSL1 knockdowns decreases PINK1-dependent activity. **A**. Representative images of pUb(Ser65) (green) following treatment of SCR, PINK1, KAT8 and KANSL1 KD POE SH-SY5Y cells with 1 μM O/A for 0-7 h. Insets show the nuclei for the same fields. Scale bar: 20 μm. **B**. Quantification of pUb(Ser65) in A (n=6, two-way ANOVA with Dunnett’s correction). For details on the statistical test, see Supplementary Table 4. **C**. Representative images of FLAG-Parkin (green) following treatment of SCR, PINK1 and KAT8 siRNA KD POE SH-SY5Y with 1 µM O/A for 3 h. Scale bar: 20 µm. **D**. Quantification of FLAG-Parkin recruitment to the mitochondria as a ratio of FLAG intensity in the mitochondria and in the cytosol in C (n=5, two-way ANOVA with Dunnett’s correction). **E**. Representative images of pParkin (green) following treatment of SCR, PINK1 and KAT8 siRNA KD POE SH-SY5Y with 1 µM O/A for 3 h. Scale bar: 20 µm. **F**. Quantification of pParkin levels in E (n=5, two-way ANOVA with Dunnett’s correction). Data are shown as mean ± SD.

**Figure 4.**
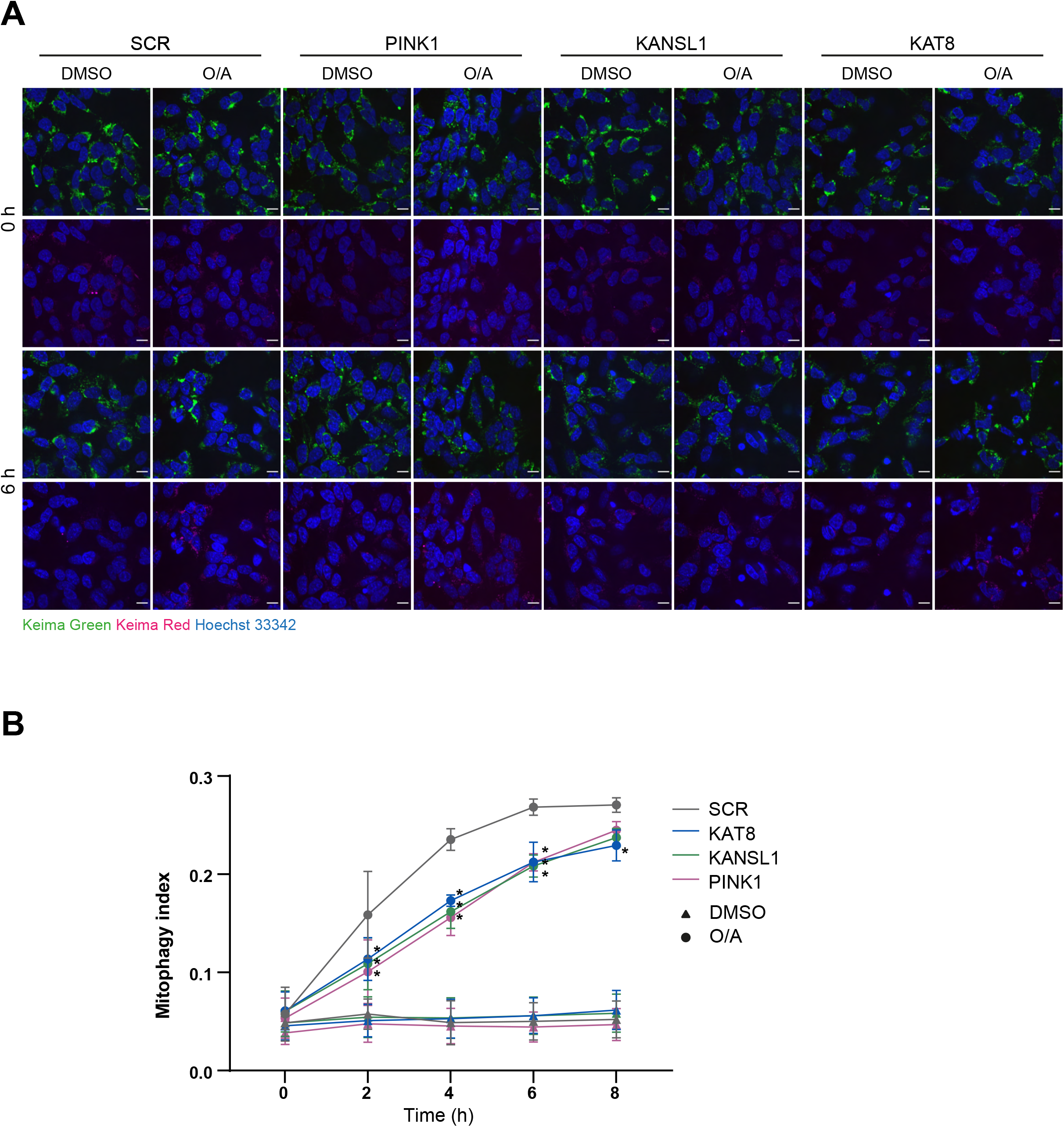
KANSL1 and KAT8 knockdown decrease mitochondrial clearance. **A**. Representative images of mt-Keima following treatment of SCR, PINK1 and KAT8 siRNA KD POE SH-SY5Y with 1 µM O/A for 0-8 h. Scale bar: 25 µm. **B**. Quantification of the mitophagy index, calculated as the ratio of the area of lysosomal mt-Keima signal and total mt-Keima signal in A (n=3, one-way ANOVA with Dunnett’s correction). For details on the statistical test, see Supplementary Table 5. Data are shown as mean ± SD.

*KANSL1* is located within the extensively studied inversion polymorphism on chromosome 17q21 (Extended Data Fig. 8A, B), which also contains *MAPT* - a gene frequently postulated to drive PD risk at this locus (Wray and Lewis, 2010). While the majority of individuals inherit this region in the direct orientation, up to 25% of individuals of European descent have a ∼ 1mb sequence in the opposite orientation (Stefansson *et al*., 2005; Zody *et al*., 2008), inducing a larger ∼ 1.3–1.6 Mb region of linkage disequilibrium (LD). Since this inversion polymorphism precludes recombination over a region of ∼ 1.3–1.6 Mb, haplotype-specific polymorphisms have arisen resulting in the generation of two major haplotype clades, termed H1 (common haplotype) and H2 (inversion carriers), previously strongly linked to neurodegenerative disease (Hutton *et al*., 1998; Pittman *et al*., 2005). Due to high LD, the genetics of this region have been hard to dissect, and robust eQTL analyses have been challenging due to the issue of polymorphisms within probe sequences in microarray-based analyses or mapping biases in RNA-seq-based analyses. Several variants (rs34579536, rs35833914 and rs34043286) are in high LD with the H1/H2 haplotype and are located within *KANSL1* (Fig. 5A,B), raising the possibility that they could directly impact on KANSL1 protein function. In particular, one of the missense variants is a serine to proline change in KANSL1 protein sequence (S718P), and would therefore be predicted to alter the gross secondary structure of the KANSL1 protein (Fig. 5B). Furthermore, we explored the possibility that PD risk might be mediated at this locus through an effect on *KANSL1* expression. Using RNA sequencing data generated from 84 brain samples (substantia nigra n=35; putamen n=49), for which we had access to whole exome sequencing and SNP genotyping data thus enabling mapping to personalised genomes (Guelfi *et al*., 2019), we performed allele-specific expression analysis. More specifically, we quantified the variation in expression between the H1 and H2 haplotypes (Supplementary Table 6) amongst heterozygotes. While we identified ASE sites within *MAPT* (Extended Data Fig. 9 and Supplementary Table 7), we also identified 4 sites of allele-specific expression in *KANSL1* (Fig. 5A), suggesting that the high PD risk H1 allele is associated with lower *KANSL1* expression, consistent with our functional assessment. Interestingly, sequence analysis of the human *KANSL1* haplotype revealed that the high risk H1 haplotype is the more recent “mutant” specific to *Homo sapiens*, and that other primates and mammals share the rarer non-risk ancestral H2 haplotype (Fig. 5B). To assess the specificity of the KANSL1 KD effect on PINK1-mitophagy, 32 open reading frames in linkage disequilibrium on the H1 haplotype at the 17q21 locus (Extended Data Fig. 8A, B and Supplementary Table 8) were knocked down individually and their effect on pUb(Ser65) was assessed. While the effect of *KANSL1* KD on pUb(Ser65) was confirmed, neither the KD of *MAPT*, nor the KD of each of the other 30 genes on this locus, led to a decreased in the pUb(Ser65) signal (Fig. 5C). These data confirm the selectivity of our mitophagy screening assay and suggest that *KANSL1* is likely to be a key PD risk gene at the 17q21 locus.

**Figure 5.**
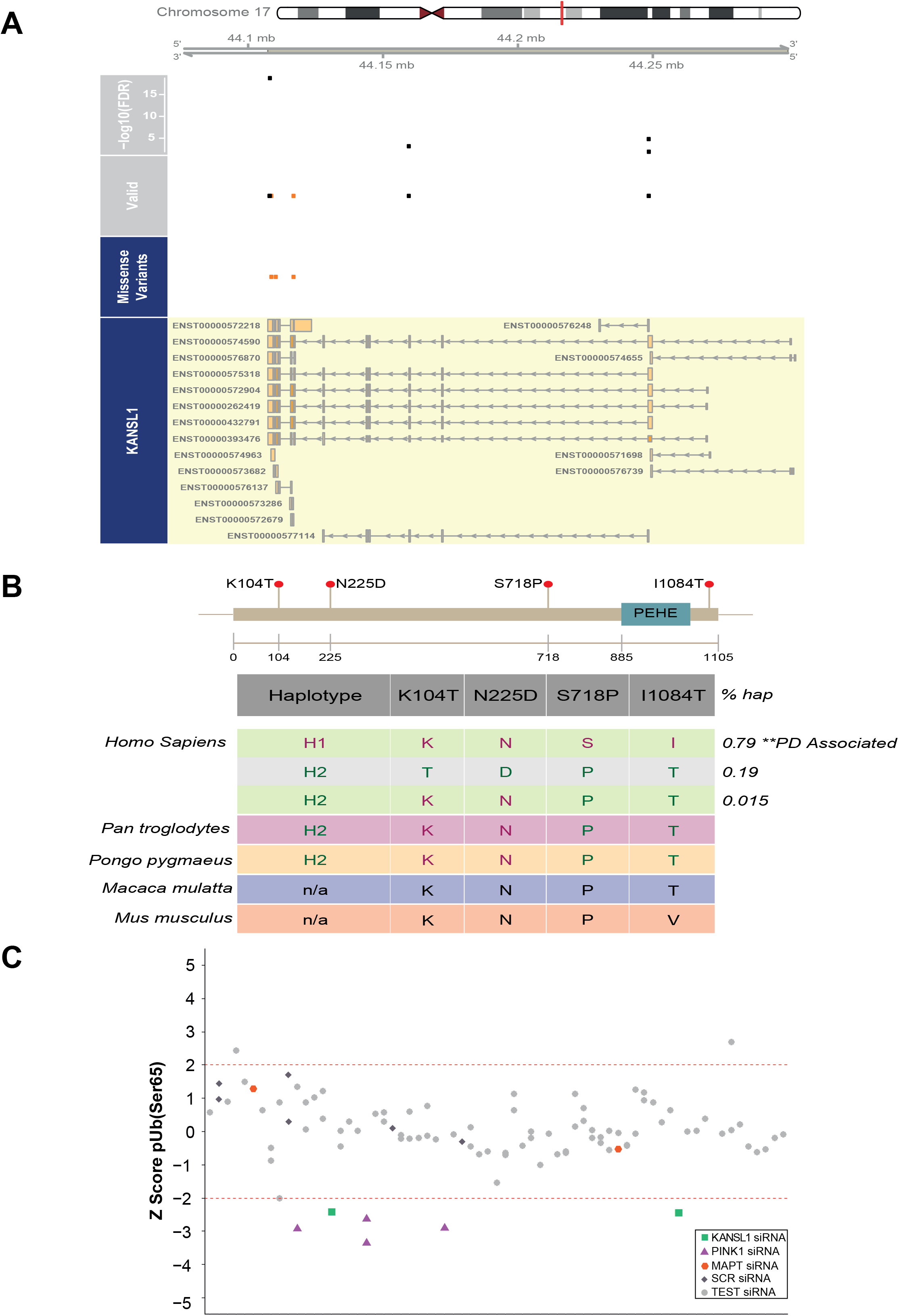
KANSL1, which presents ASE sites in LD with the H1/H2 SNP, is the only gene on the 17q21 locus modulating pUb(Ser65) levels. **A**. ASEs derived from putamen and substantia nigra in high linkage disequilibrium with the H1/H2 tagging SNP, rs12185268 and their position along the *KANSL1* gene. The missense variants track displays the variants annotated as missense by gnomAD v2.1.1(Lek *et al*., 2016). The valid track displays the heterozygous sites (orange = missense) with an average read depth greater than 15 reads across all samples, which were examined for ASE. The topmost track displays the FDR-corrected minimum -log10 p-value across samples for the sites that show an ASE in at least one sample. **B**. Conservation of the KANSL1 protein across species. The four coding variants in the *KANSL1* gene are in high LD (r2 >0.8) with the H1/H2 haplotypes. **C**. pUb(Ser65) Z-scores of one representative 17q21 locus screen plate. See Supplementary Table 8 for the complete list of the genes screened.

Finally, we sought to study the mechanism of disrupted mitophagy in KAT8 and KANSL1 deficient cells. KAT8 and the NSL complex are mainly responsible for the acetylation of lysine 16 on histone 4, and are therefore master regulators of transcription (Sheikh, Guhathakurta and Akhtar, 2019). As a result, we hypothesised that they may regulate PINK1-mitophagy by regulating *PINK1* gene transcription. In order to test that hypothesis, we knocked down KAT8 and KANSL1 in POE SHSY5Y cells before extracting RNA and performing qPCR. KD of KANSL1 significantly reduced PINK1 mRNA levels (Fig. 6B), while KAT8 had a modest effect, suggesting that KANSL1 KD may affect PINK1-mitophagy by modulating *PINK1* mRNA levels.

**Figure 6.**
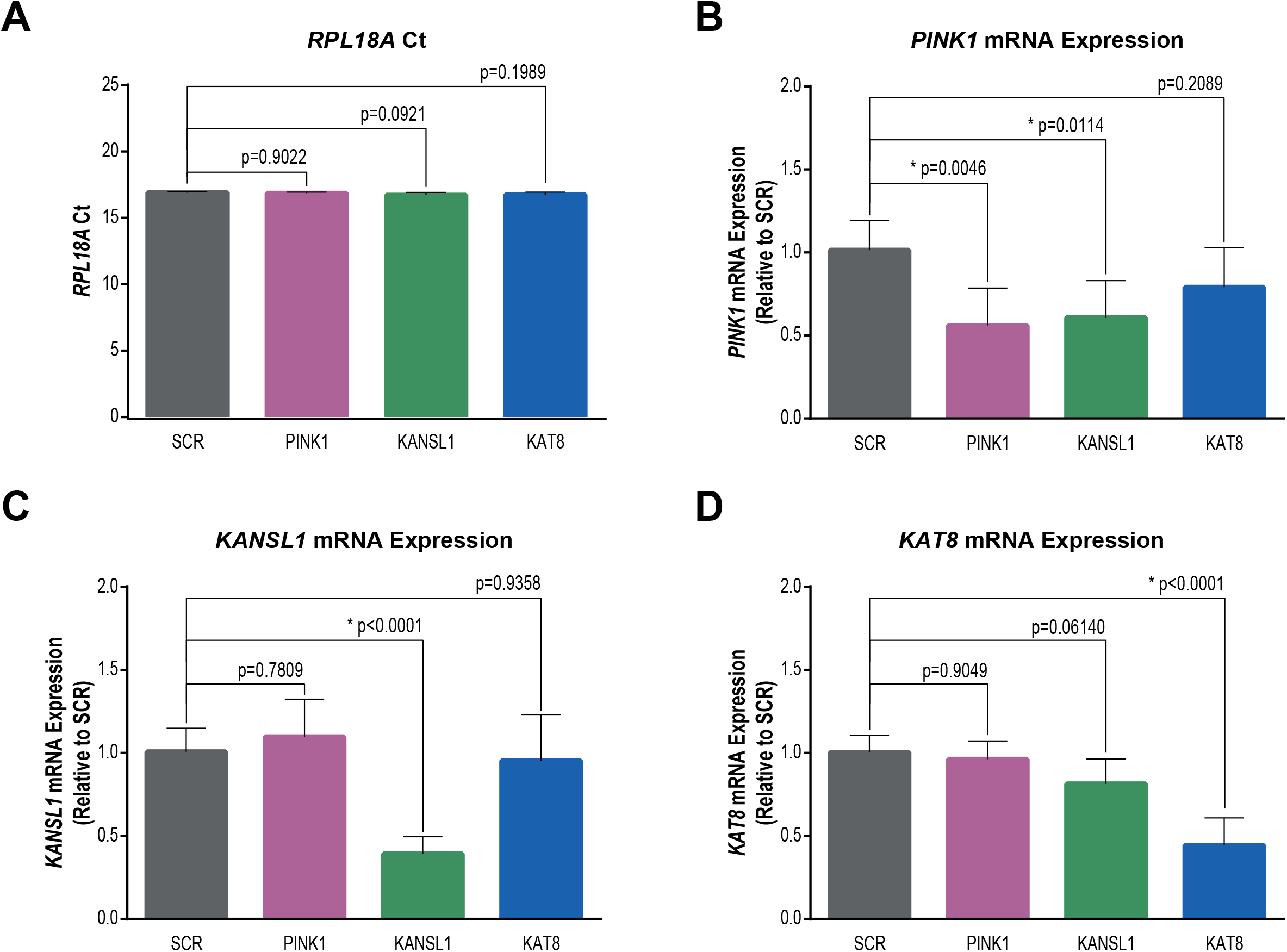
KANSL1 knockdown reduces PINK1 mRNA levels in POE SHSY5Y cells. **A**. Ct values for *RPL18A* were unaffected by siRNA KD. n=6, one-way ANOVA with Dunnett’s correction. **B**. Relative *PINK1* mRNA expression levels in SCR, PINK1, KANSL1 and KAT8 siRNA KD POE SH-SY5Ys, as measured through RT-qPCR (n=6, one-way ANOVA with Dunnett’s correction). **C**. Relative *KANSL1* mRNA expression levels in SCR, PINK1, KANSL1 and KAT8 siRNA KD POE SH-SY5Ys,as measured through RT-qPCR (n=6, one-way ANOVA with Dunnett’s correction). **D**. Relative *KAT8* mRNA expression levels in SCR, PINK1, KANSL1 and KAT8 siRNA KD POE SH-SY5Ys, as measured through RT-qPCR (n=6, one-way ANOVA with Dunnett’s correction). Data are shown as mean ± SD.

## DISCUSSION

Since the first PD GWAS study was performed in 2006 (Fung *et al*., 2006), GWAS have identified about 90 independent loci for PD (Nalls *et al*., 2019). However, translating GWAS findings into a new molecular understanding of PD-associated pathways and new therapeutic targets has remained a major challenge for the scientific community. In order to screen for PD GWAS candidate genes that play a role in PINK1-mitophagy, and thus are likely to be genuine risk genes for PD, we have set up and optimised a HCS for pUb(Ser65), a marker of PINK1-dependent mitophagy, a key pathway in PD pathogenesis. This approach allowed the successful identification of two new genes associated with increased PD risk, that play a role in mitophagy. Interestingly, these two genes were previously shown to be part of the same complex, the NSL complex.

This study demonstrates the substantial potential of functional screens to exploit genetic data by providing orthogonal information that can confidently identify new risk genes. This is particularly important in genomic regions with uniformly high linkage disequilibrium, such as the 17q21 inversion region which includes 32 ORFs of which many are highly expressed in brain and where existing fine-mapping and functional genomic analyses have been inconclusive. Interestingly, while *MAPT* has long been considered the risk associated gene at this locus, this has recently been questioned by Dong and colleagues, who also raised the significance of KANSL1 in driving PD risk at the locus (Dong *et al*., 2018). Furthermore, functional screening can simultaneously provide mechanistic insights as exemplified in this case by the novel insights we provide into the molecular events regulating mitochondrial quality control and which support a role for mitophagy as a contributing factor to sporadic PD. The KAT8 and KANSL1-containing NSL complex was shown to promote histone acetylation and as such, is a master regulator of transcription (Sheikh, Guhathakurta and Akhtar, 2019). Our data demonstrate that KAT8 and KANSL1 modulate *PINK1* nuclear transcription and subsequent translation, leading to regulation of PINK1-dependent mitophagy. It was previsouly shown that depletion of KAT8/KANSL1 causes significant downregulation of mitochondrial DNA transcription and translation, and ultimately impaired mitochondrial respiration (Chatterjee *et al*., 2016). Future studies will need to determine whether KAT8/KANSL1-dependent modulation of mitochondrial DNA could regulate PINK1 mitochondrial accumulation and subsequent mitophagy. It has been further proposed that the KAT8/KANSL1 complex has targets in the mitochondria other than the mitochondrial DNA (Chatterjee *et al*., 2016). It will be interesting to determine whether the KAT8/KANSL1 complex could acetylate ubiquitin, which has previously been shown to be acetylated on six out of its seven lysines (K6, K11, K27, K33, K48, K63) (Swatek and Komander, 2016).

Important genetic discoveries in PD, in particular, the identification of the *PINK1* (Valente *et al*., 2004) and *PRKN* genes (Kitada *et al*., 1998), opened the field of selective mitophagy (McWilliams and Muqit, 2017). However, there is still a clear need for a better molecular understanding of mitochondrial quality control. Here we provide new insights into the mechanism by identifying two new molecular players, KAT8 and KANSL1. These new regulators of mitophagy provide the first direct evidence for a role of the PINK1-mitophagy pathway in idiopathic PD and the convergence between familial and idiopathic pathways in disease. Taken together, these findings open a novel avenue for the therapeutic modulation of mitophagy in PD, with potential implications across drug discovery in frontotemporal dementia and Alzheimer’s disease, where mitophagy also plays an important role in disease pathogenesis (Chu, 2019).

## METHODS

### Reagents

Oligomycin (mitochondrial complex V inhibitor) was purchased from Cayman Chemicals (11341) and from Sigma-Aldrich (O4876), and antimycin A (mitochondrial complex III inhibitor) was purchased from Sigma-Aldrich (A8674). All siRNAs were purchased as pre-designed siGENOME SMARTpools from Dharmacon: non-targeting (D-001206-13), PINK1 (M-004030-02), PLK1 (L-003290-00), KIF-11 (L-003317-00), KAT8 (M-014800-00), KANSL1 (M-031748-00), KANSL2 (M-020816-01), KANSL3 (M-016928-01), HCFC1 (M-019953-01), MCRS1 (M-018557-00), OGT (M-019111-00), PHF20 (M-015234-02), WDR5 (M-013383-01). The following antibodies were used for immunocytochemistry: mouse anti TOM20 (Santa Cruz, sc-17764, 1:1000), rabbit anti phospho-ubiquitin (Ser65) (Cell Signaling, 37642, 1:1000), rabbit anti phospho-Parkin (Ser65) (Abcam/Michael J. Fox Foundation, MJF17, 1:250), rabbit anti FLAG (Sigma-Aldrich, F7425, 1:500),, AlexaFluor 488 goat anti rabbit (Invitrogen, A11008, 1:2000), AlexaFluor 568 goat anti mouse (Invitrogen, A11004, 1:2000),. The following antibodies were used for immunoblotting: mouse anti TIM23 (BD Biosciences, 611223, 1:1000), rabbit anti phospho-ubiquitin (Ser65) (Merck Millipore, ABS1513-I, 1:1000), mouse anti GAPDH (Abcam, ab110305, 1:1000), rabbit anti KAT8 (Abcam, ab200600, 1:1000), IRDye 680LT donkey anti mouse (LI-COR Biosciences, 925-68022, 1:20000), IRDye 800CW donkey anti rabbit (LI-COR Biosciences, 925-32213, 1:20000).

### Selection of genes for High Content Screening

Candidates for High Content Screening were selected based on i) WPPINA; ii) complex prioritization; and, iii) coloc analysis. WPPINA analysis is reported in (Ferrari *et al*., 2018) where the 2014 PD GWAS (Nalls *et al*., 2014) was analysed; candidate genes where selected among those prioritised and with an LD r2 ≥ 0.8. The same pipeline has then been additionally applied to the 2017 PD GWAS (Chang *et al*., 2017) to update the list of candidate genes. Briefly, a protein-protein interaction network has been created based on the Mendelian genes for PD (seeds) using data from databases within the IMEx consortium. The network has been topologically analysed to extract the core network (i.e. the most interconnected part of the network). The core network contains the proteins/genes that can connect >60% of the initial seeds and are therefore considered relevant for sustaining communal processes and pathways, shared by the seeds. These processes have been evaluated by Gene Ontology Biological Processes enrichment analysis. The top SNPs of the 2017 PD GWAS have been used to extract open reading frames (ORFs) in cis-haplotypes defined by LD r2 ≥ 0.8 (analysis performed in October 2017). These ORFs have been matched to the core network to identify overlapping proteins/genes in relevant/shared pathways. Results of complex prioritization (neurocentric prioritization strategy) were gathered from (Chang *et al*., 2017) where this strategy was applied to the 2017 PD GWAS. The coloc analysis was performed as reported in (Kia *et al*., 2019), posterior probabilities for the hypothesis that both traits, the regulation of expression of a given gene and the risk for PD share a causal variant (PPH4), were calculated for each gene, and genes with PPH4 ≥ 0.75 were considered to have strong evidence for colocalization. Summary statistics were obtained from the most recent PD GWAS (Nalls *et al*., 2019) and were used for regional association plotting using LocusZoom (Pruim *et al*., 2010).

### Cell Culture and siRNA transfection

POE SH-SY5Y cells are a kind gift from H. Ardley (Ardley *et al*., 2003) and the mt-Keima POE SHSY5Y cells were a kind gift of C. Luft (Soutar *et al*., 2019). Cells were cultured in Dulbecco’s Modified Eagle (DMEM, Gibco, 11995-065) and supplemented with 10% heat-inactivated foetal bovine serum (FBS, Gibco) in a humidified chamber at 37 °C with 5% CO_2_. For siRNA transfection, cells were transfected using DharmaFECT1 transfection reagent (Dharmacon, T-2001-03) according to the manufacturer’s instructions (for concentrations of siRNA, see sections below).

### ASEs

Sites of ASE were identified as described by Guelfi and colleagues (Guelfi *et al*., 2019) by mapping RNA-seq data to personalised genomes, an approach specifically chosen because it aims to minimise the impact of mapping biases. RNA-seq data generated from 49 putamen and 35 substantia nigra tissue samples from the UK Brain Expression Consortium was used for this analysis. All samples were obtained from neuropathologically normal individuals of European descent and sites with greater than 15 reads in a sample were tested for ASE. Only sites present in non-overlapping genes were considered and data from both the tissues were considered together to increase power. Sites with minimum FDR < 5% across samples were marked as ASE sites. Plots were generated using Gviz3, with gene and transcript details obtained from Ensembl v92.

### High Content siRNA Screen

#### Cell plating and siRNA transfection

siRNA was dispensed into Geltrex-coated 96-well CellCarrier Ultra plates (Perkin Elmer) at a final concentration of 30 nM using the Echo 555 acoustic liquid handler (Labcyte). For each well, 25 µl of DMEM containing 4.8 µl/ml of DharmaFECT1 transfection reagent was added and incubated for 30 min before POE SH-SY5Y cells were seeded using the CyBio SELMA (Analytik Jena) at 15,000 cells per well, 100 µl per well in DMEM + 10% FBS. Cells were incubated for 72 h before treatment with 10 µM oligomycin/10 µM antimycin for 3 h to induce mitophagy.

#### IF and Image Capture and Analysis

Cells were fixed with 4% PFA (Sigma-Aldrich, F8775), then blocked and permeabilised with 10% FBS, 0.25% Triton X-100 in PBS for 1 h, before immunostaining with pUb(Ser65) and TOM20 primary antibodies (in 10% FBS/PBS) for 2 h at room temperature. After 3x PBS washes, AlexaFluor 568 anti-mouse and 488 anti-rabbit secondary antibodies and Hoechst 33342 (Thermo Scientific, 62249) were added (in 10% FBS/PBS, 1:2000 dilution for all) and incubated for 1 h at room temperature. Following a final 3x PBS washes, plates were imaged using the Opera Phenix (Perkin Elmer). 5x fields of view and 4x 1 µm Z-planes were acquired per well, using the 40X water objective, NA1.1. Images were analysed in an automated way using the Columbus 2.8 analysis system (Perkin Elmer) to measure the integrated intensity of pUb(Ser65) within the whole cell. First of all, the image was loaded as a maximum projection, before being segmented to identify the nuclei using the Hoechst 33342 channel (method B). The cytoplasm was then identified using the “Find Cytoplasm” building block (method B) on the sum of the Hoechst and Alexa 568 channels. pUb(Ser65) was identified as spots (method B) on the Alexa 488 channel, before measuring their integrated intensity.

#### Screen quality control, data processing and candidate selection

Screen plates were quality controlled based on the efficacy of the PINK1 siRNA control and O/A treatment window (minimum 3-fold). Data were checked for edge effects using Dotmatics Vortex visualization software. Raw data was quality controlled using robust Z prime > 0.5. Data were processed using Python for Z score calculation before visualization in Dotmatics Vortex. Candidates were considered a hit where Z score was ≥ 2 or ≤ -2, and where replication of efficacy was seen both within and across plates.

#### siRNA libraries

The siRNA libraries were purchased from Dharmacon as an ON-TARGETplus SMARTpool Cherry-pick siRNA library, 0.25 nmol in a 384-well plate. siRNAs were resuspended in RNase-free water for a final concentration of 20 µM. SCR, PINK1 and PLK1 or KIF11 controls were added to the 384-well plate at a concentration of 20 µM before dispensing into barcoded assay-ready plates.

### Mitochondrial enrichment and Western blotting and WES

POE SH-SY5Y and H4 cells were transfected with 100 nM siRNA and incubated for 72 h. Whole cell lysates were used from H4 cells, whereas POE SH-SY5Y lysates were first fractionated into cytoplasmic and mitochondria-enriched preparations. Samples were run on SDS-PAGE before IB with the Odyssey® CLx Imager (LI-COR Biosciences). Mitochondrial enrichment and Western blotting protocols were described previously (Soutar *et al*., 2018).

### Immunofluorescence

POE SH-SY5Y cells were reverse transfected with 50 nM siRNA in 96-well CellCarrier Ultra plates according to the manufacturer’s instructions and incubated for 72 h. Cells were then treated, fixed and stained as per the screening protocol detailed above (for treatment concentrations and times, see figures). For visualisation purposes, brightness and contrast settings were selected on the SCR controls and applied to all other images. Images are presented as maximum projections of the channels for one field of view. Insets show the Hoechst 33342 channel for the same field.

### Mitophagy measurement using the mt-Keima reporter

Stable mt-Keima expressing POE SHSY5Y cells were reverse transfected with 50 nM siRNA in 96-well CellCarrier Ultra plates according to the manufacturer’s instructions and incubated for 72 h. For the assay, the cell medium was replaced with phenol-free DMEM + 10% FBS containing Hoechst 33342 (1:10000) and either DMSO or 1 µM oligomycin/1 µM antimycin to induce mitophagy. Cells were immediately imaged on the Opera Phenix (PerkinElmer) at 37 °C with 5% CO_2_, acquiring 15x single plane fields of view, using the 63X water objective, NA1.15. The following excitation wavelengths and emission filters were used: cytoplasmic Keima: 488 nm, 650–760 nm; lysosomal Keima: 561 nm, 570–630 nm; Hoechst 33342: 375 nm, 435–480 nm. Images were analysed in an automated way using the Columbus 2.8 analysis system (Perkin Elmer) to measure the mitophagy index. Cells were identified using the nuclear signal of the Hoechst 33342 channel, before segmenting and measuring the area of the cytoplasmic and lysosomal mt-Keima. The mitophagy index was calculated as the ratio between the total area of lysosomal mitochondria and the total area of mt-Keima (sum of the cytoplasmic and lysosomal mtKeima areas) per well.

### RT-qPCR

Total RNA was extracted from cells using the Monarch Total RNA Miniprep Kit (New England Bioscience) with inclusion of the optional on-column DNAse treatment. 500ng of the RNA was then reverse transcribed with SuperScript^™^ IV reverse transcriptase (Invitrogen). The cDNA product was then subjected to quantitative real-time PCR (qPCR) using Fast SYBR^™^ Green Master Mix (Applied Biosystems) and gene specific primer pairs (Supplementary Table 9) on a QuantStudio^™^ 7 Flex Real-Time PCR System (Applied Biosystems). Relative mRNA expression levels were calculated using the 2^−ΔΔCt^ method and *RPL18A* as the house-keeping gene.

### *Drosophila* stocks and husbandry

Flies were raised under standard conditions in a humidified, temperature-controlled incubator with a 12h:12h light:dark cycle at 25°C, on food consisting of agar, cornmeal, molasses, propionic acid and yeast. The following strains were obtained from the Bloomington *Drosophila* Stock Center (RRID:SCR_006457): *mof* RNAi lines, P{TRiP.JF01701} (RRID:BDSC_31401); and P{TRiP.HMS00537} (RRID:BDSC_58281); *nsl1* RNAi lines, P{TRiP.HMJ22458} (RRID:BDSC_58328); the pan-neuronal *nSyb-GAL4* driver (RRID:BDSC_51941); and dopaminergic neuron driver (TH-GAL4; RRID:BDSC_8848); and control (*lacZ*) RNAi P{GD936}v51446) from the Vienna *Drosophila* Resource Center (RRID:SCR_013805). All experiments were conducted using male flies.

### Locomotor and lifespan assays

The startle induced negative geotaxis (climbing) assay was performed using a counter-current apparatus. Briefly, 20-23 males were placed into the first chamber, tapped to the bottom, and given 10 s to climb a 10 cm distance. This procedure was repeated five times (five chambers), and the number of flies that has remained into each chamber counted. The weighted performance of several group of flies for each genotype was normalized to the maximum possible score and expressed as *Climbing index* (Greene *et al*., 2003).

For lifespan experiments, flies were grown under identical conditions at low-density. Progeny were collected under very light anaesthesia and kept in tubes of approximately 20 males each, around 50-100 in total. Flies were transferred every 2-3 days to fresh medium and the number of dead flies recorded. Percent survival was calculated at the end of the experiment after correcting for any accidental loss.

### Immunohistochemistry and sample preparation

*Drosophila* brains were dissected from aged flies and immunostained as described previously (Whitworth *et al*., 2005). Adult brains were dissected in PBS and fixed in 4% formaldehyde for 30 min on ice, permeabilized in 0.3% Triton X-100 for 3 times 20 min, and blocked with 0.3% Triton X-100 plus 4% goat serum in PBS for 4 h at RT. Tissues were incubated with anti-tyrosine hydroxylase (Immunostar Inc. #22491), diluted in 0.3% Triton X-100 plus 4% goat serum in PBS for 72 h at 4°C, then rinsed 3 times 20 min with 0.3% Triton X-100 in PBS, and incubated with the appropriate fluorescent secondary antibodies overnight at 4°C. The tissues were washed 2 times in PBS and mounted on slides using Prolong Diamond Antifade mounting medium (Thermo Fisher Scientific). Brains were imaged with a Zeiss LMS 880 confocal. Tyrosine hydroxylase-positive neurons were counted under blinded conditions.

### Statistical Analysis

Intensity measurements from imaging experiments were normalised to SCR O/A for each experiment and presented as a percentage. N numbers are shown in figure legends and refer to the number of independent, replicate experiments. Within each experiment, the mean values of every condition were calculated from a minimum of 3 technical replicates. Intensity measurements from Western blot experiments were normalised to PINK1 O/A. GraphPad Prism 9 (La Jolla, California, USA) was used for statistical analyses and graph production. Data were subjected to either one-way or two-way ANOVA with Dunnett’s post-hoc analysis for multiple comparisons, unless otherwise stated. All error bars indicate mean ± standard deviation (SD) from replicate experiments.

## Acknowledgements

This work was supported in part by the UK Medical Research Council (MRC) funding to the Dementia Platform UK (MR/M02492X/1), MRC core funding to the High-Content Biology Platform at the MRC-UCL LMCB university unit (MC_U12266B) and MRC MBU (MC_UU_00015/6), and by UCL Translational Research Office administered seed funds. MS, EA, CM and DT are funded by MRC MR/N026004/1. DM is supported by an MRC CASE studentship (MR/P016677/1). BO is supported by the Michael J. Fox Foundation for Parkinson’s Research (MJFF-010437). AM, MB and PW are funded by ARUK (ARUK-2018DDI-UCL). MR was supported by the UK MRC through the award of Tenure-track Clinician Scientist Fellowship (MR/N008324/1). This work was supported in part by the Intramural Research Programs of the National Institute on Aging (NIA). We also acknowledge the support of the NIHR BRC award to University College London Hospitals, UCL. Finally, the authors would like to thank the Genome Aggregation Database (gnomAD) and the groups that provided exome and genome variant data to these resources. A full list of contributing groups can be found at https://gnomad.broadinstitute.org/about.

## Author Contributions

HPF, PL, JH, AW and PW conceived the idea. MS, DM, BO, EA, AM, DT, MB, PW, JH, AW, MR, PL and HPF designed the experiments. MS, DM, BO, EA, AM, NW, NW, KDS, SG, DZ, AP, DT, KP, CM, CB and HPF carried out analysis and experiments. MS, DM, BO, EA, AM, PW, CM, AW, MR, PL and HPF wrote the manuscript, with input from all co-authors. HPF, PL and MR supervised the project.

## EXTENDED FIGURE LEGENDS

**Extended Data Figure 1. High Content siRNA Screen for modulators of pUb(Ser65)**.

**A**. Venn diagram highlighting the three genes prioritised by means of three prediction techniques.

**B**. Fold decrease in TOM20 levels following 1.5 and 3 h treatment with 0.1, 1 and 10 µM O/A, compared to DMSO control.

**C**. Representative images of TOM20 and pUb(Ser65) following 3 h treatment of SCR KD POE SH-SY5Y cells with 10 µM O/A. Scale bar: 20 µm.

**D**. Quantification of the co-localization in **C** as % of TOM20-positive pUb(Ser65) spots. Graph shows all replicates of non-transfected, SCR, PINK1 and PLK1 KD for 3 independent experiments.

**E**. Representative images of pUb(Ser65) following treatment of SCR and PINK1 KD POE SH-SY5Y cells with 10 µM O/A for 3 h. Scale bar: 20 µm.

**F**. Quantification of pUb(Ser65) in **E** (n=6, two-way ANOVA with Tukey’s multiple comparisons test).

**G**. Representative analysis of integrated intensity of pUb(Ser65) and TOM20 for a single HCS plate.

**H**. pUb(Ser65) Z-scores of the two other replicate screen plates.

Data are shown as mean ± SD.

**Extended Data Figure 2. KAT8 knockdown has no effect on cell viability**.

**A**. Representative images of nuclei following treatment of SCR, PINK1 and PLK1 siRNA KD POE SH-SY5Y cells with 10 µM O/A for 3 h. Scale bar: 20 µm.

**B**. Quantification of the number of nuclei in A (n=6, two-way ANOVA with Tukey’s multiple comparisons test).

**C**. Z-scores of a representative screen plate showing that KAT8 or PINK1 siRNA KD don’t affect cell viability, on the contrary to PLK-1 KD. Data are shown as mean ± SD.

**Extended Data Figure 3. KAT8 eQTLs colocalise with SNPs associated with PD risk**

The x-axis displays the physical position on chromosome 16 in megabases. The minus log p-values are plotted for every SNP present in both the PD GWAS (Chang *et al*., 2017) and *KAT8* eQTLs derived from the GTEx V7 caudate data. The p-values for the PD GWAS are plotted in yellow and p-values for *KAT8* eQTLs are plotted in blue.

**Extended Data Figure 4. KAT8 knockdown decreases pUb(Ser65) levels**.

**A**. Representative images of pUb(Ser65) following treatment of SCR, PINK1 and KAT8 siRNA KD POE SH-SY5Y with 1 µM O/A for 3 h. Insets show nuclear staining for the same fields. Scale bar: 20 µm.

**B**. Quantification of pUb(Ser65) levels in A (n=3, two-way ANOVA with Dunnett’s correction).

Data are shown as mean ± SD.

**Extended Data Figure 5. Knockdown of the mitochondrial components of the NSL complex reduces pUb(Ser65) levels**.

Quantification of pUb(Ser65) following treatment of SCR, PINK1 or NSL components siRNA KD POE SH-SY5Y cells with 1 μM O/A for 3 h. Data are shown as mean ± SD; n=6, one-way ANOVA with Dunnett’s correction.

**Extended Data Figure 6. KAT8 and KANSL1 knockdown reduce pUb(Ser65) levels in WT SHSY5Y and H4 cells**.

**A**. Representative images of pUb(Ser65) following treatment of SCR, PINK1 and KAT8 siRNA KD WT SH-SY5Y with 1 µM O/A for 3 h. Insets show nuclear staining for the same fields. Scale bar: 20 µm.

**B**. Quantification of pUb(Ser65) levels in A (n=6, two-way ANOVA with Dunnett’s correction).

**C**. Representative IB of whole-cell lysates from SCR, PINK1, KANSL1 and KAT8 siRNA KD H4 cells treated with 1 μM O/A for 3 h.

**D**. Quantification of pUb(Ser65) in D (n=3, one-way ANOVA with Dunnett’s correction).

Data are shown as mean ± SD.

**Extended Data Figure 7. Neuronal loss of *mof* or *nsl1* causes locomotor deficit, shortened lifespan and neurodegeneration**.

**A, B**. Climbing ability of pan-neuronal (*nSyb-GAL4*) driven knockdown of *mof* (**A**) or *nsl1* (**B**) measured at the indicated age of adults, compared to control RNAi (A: Kruskal-Wallis test, with Dunn’s post-hoc multiple comparisons; B: Mann-Whitney test).

**C, D**. Lifespan of ***mof* (C) or *nsl1* (D)** pan-neuronal knockdown (*nSyb-GAL4*) compared to control RNAi (Log-rank (Mantel-Cox) test).

**E, F**. Quantification of dopaminergic neurons (PPL1 cluster) after pan-neuronal or dopaminergic (DA) neuron (*TH-GAL4*) driven depletion of *mof* (**E**), nsl1 (**F**), or control RNAi. Representative images of PPL1 neurons (as bounded by the box) under depletion conditions are shown. Flies were aged 30 days, except for pan-neuronal *nsl1* kd which are 16-days-old. Scale bar: 20 µm; Mann-Whitney test. For all tests, n numbers are indicated in the graphs; p<0.0001 = ****; p<0.001 = ***.

**Extended Data Figure 8. Overview of the PD GWAS genetic signal at the *MAPT* locus**.

**A**. *MAPT* primary GWAS signal.

**B**. *MAPT* conditional GWAS signal.

**Extended Data Figure 9. ASE sites in *MAPT* in LD with the H1/H2 SNP**.

ASEs derived from putamen and substantia nigra that are in LD with the H1/H2 tagging SNP, rs12185268 and their position along the *MAPT* gene. The missense variants track displays the variants annotated as missense by gnomAD v2.1.1 (Lek *et al*., 2016). The valid track displays the heterozygous sites (orange = missense) with an average read depth greater than 15 reads across all samples, in LD with H1/H2, which were examined for ASE. The topmost track displays the –log10 scale for the minimum FDR across samples for the sites that show an ASE in at least one sample.

