## Supplementary figures and images for "Regulation of mitophagy by the NSL complex underlies genetic risk for Parkinson’s disease at Chr16q11.2 and on the MAPT H1 allele"

### Extended Data Fig. 1

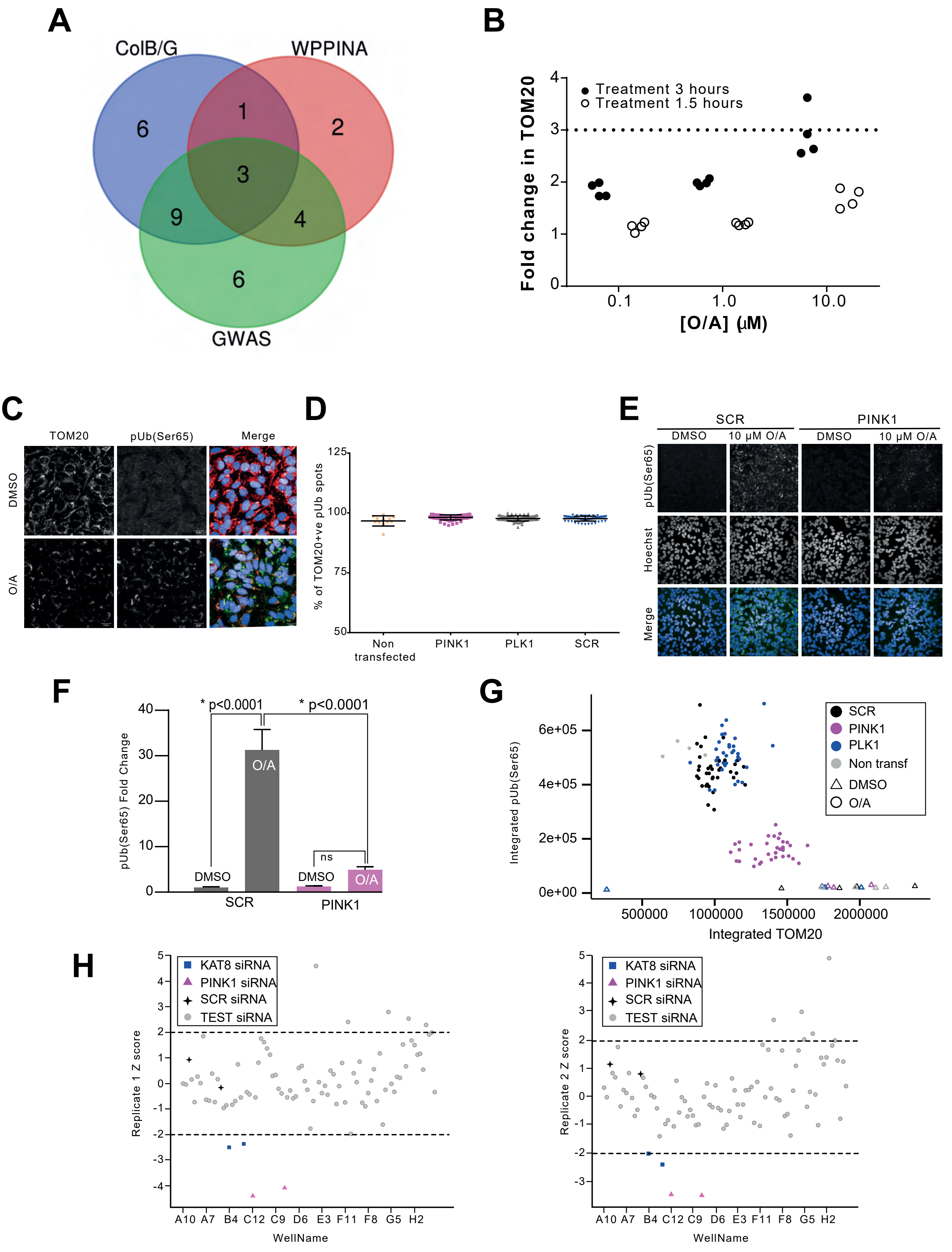


**Extended Data Figure 1 - High Content siRNA Screen for modulators of pUb(Ser65)**

### Extended Data Fig. 2

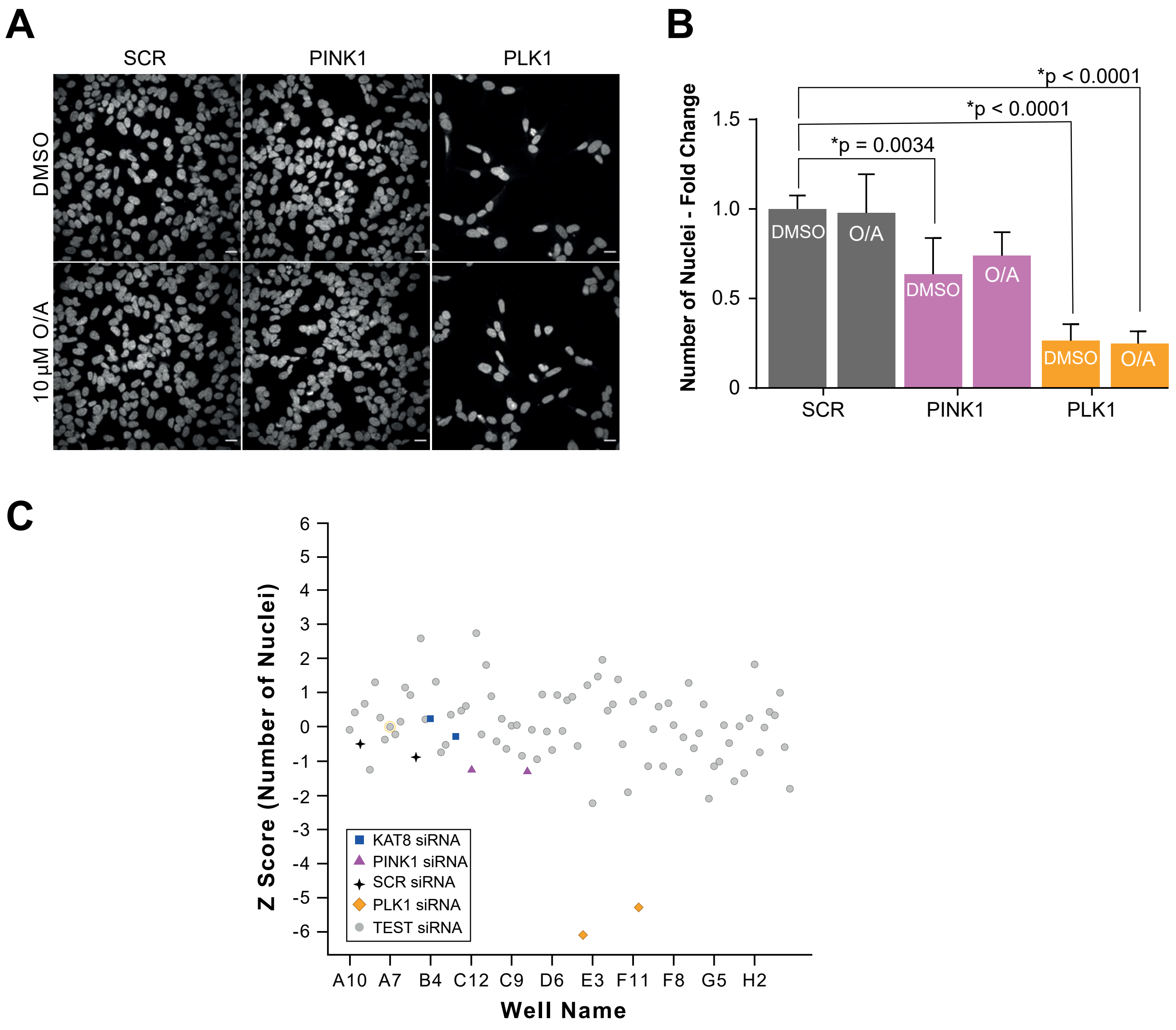


**Extended Data Figure 2 - KAT8 knockdown has no effect on cell viability**

### Extended Data Fig. 3

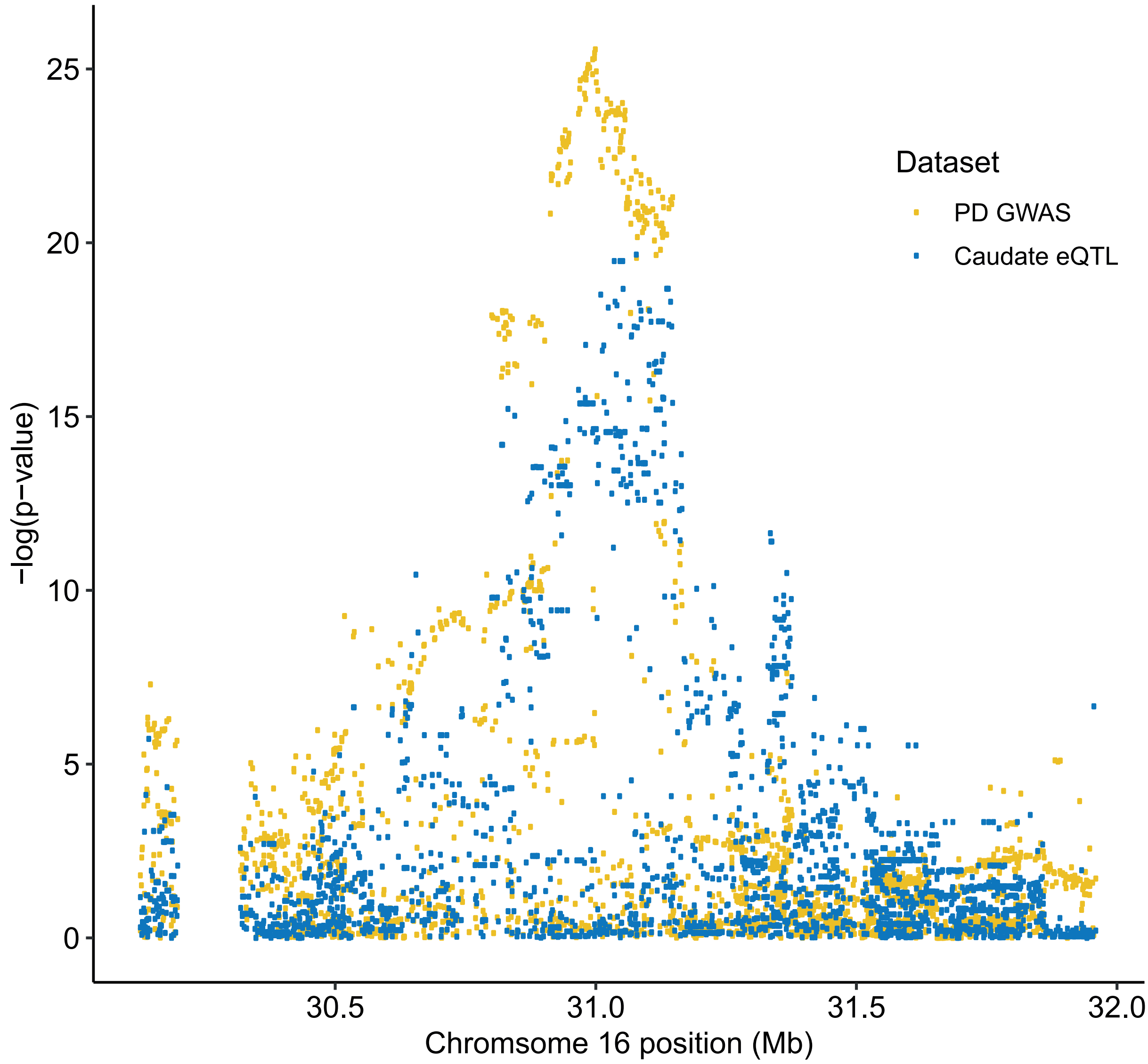


**Extended Data Figure 3 - KAT8 eQTLs colocalise with SNPs associated with PD risk**

### Extended Data Fig. 4

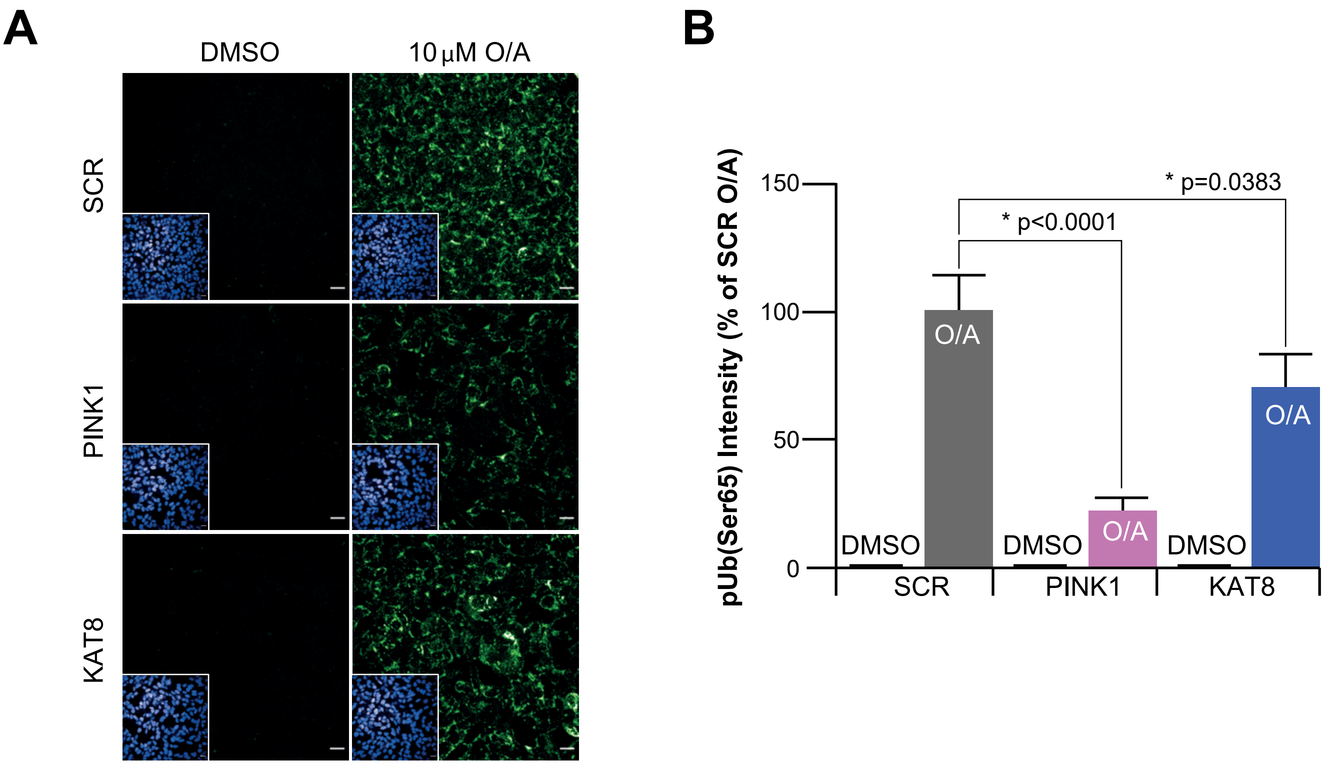


**Extended Data Figure 4 - KAT8 knockdown decreases pUb(Ser65) levels**

### Extended Data Fig. 6

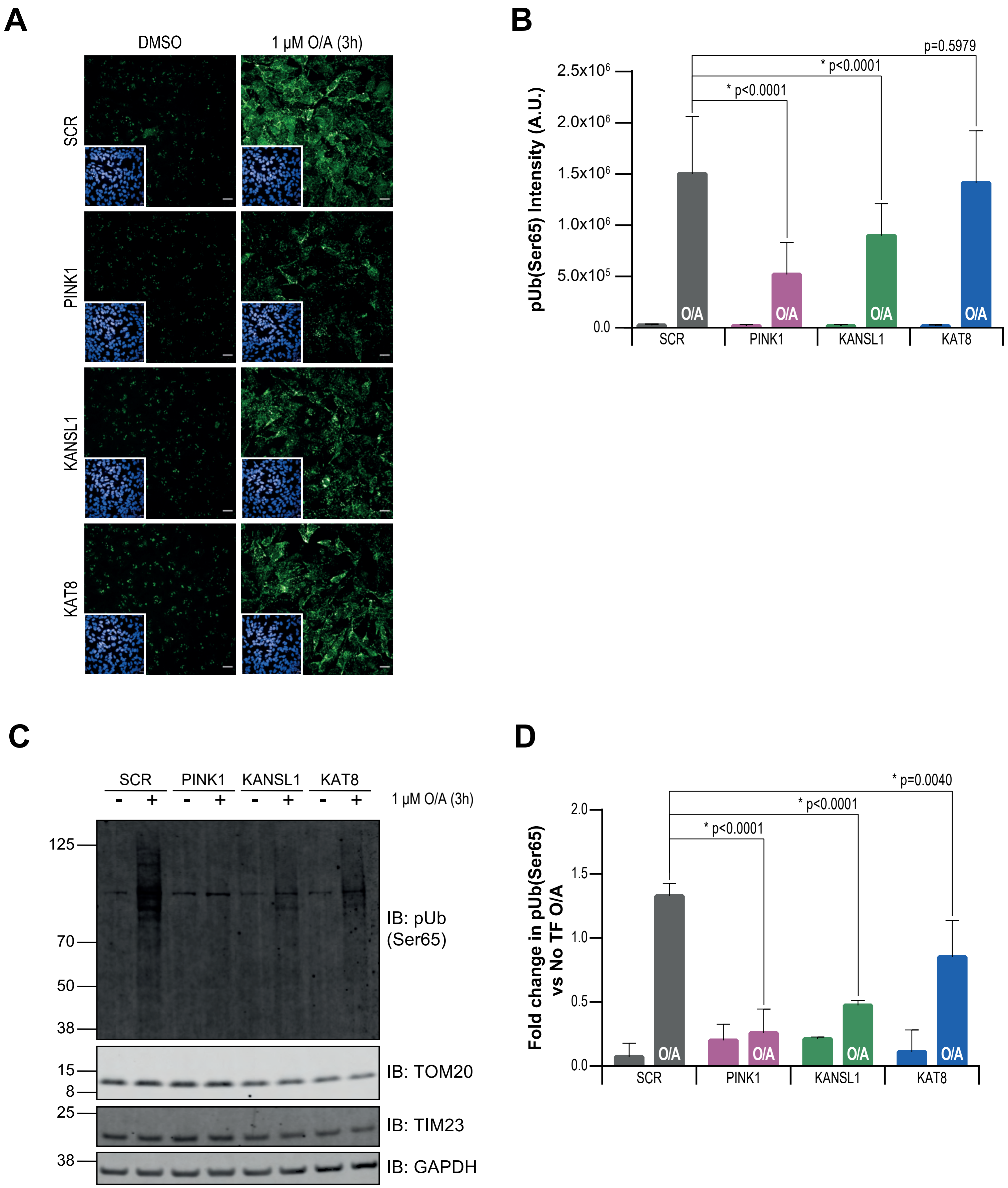


**Extended Data Figure 6 - KAT8 and KANSL1 knockdown reduce pUb(Ser65) levels in WT SHSY5Y and H4 cells**

### Extended Data Fig. 8

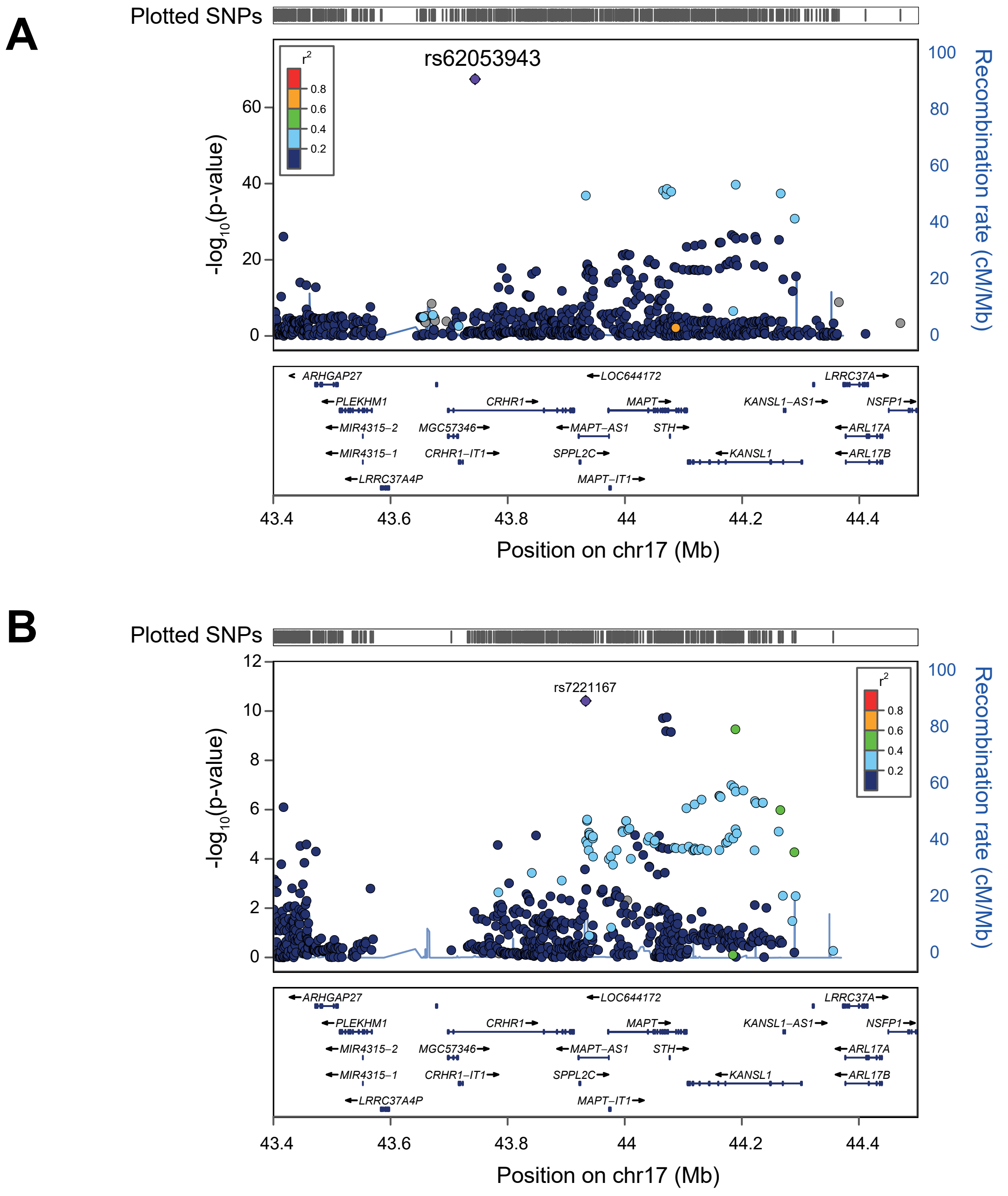


**Extended Data Figure 8 - Overview of the PD GWAS genetic signal at the *MAPT* locus**
