## Extended Data Fig. 5 for "Regulation of mitophagy by the NSL complex underlies genetic risk for Parkinson’s disease at Chr16q11.2 and on the MAPT H1 allele"

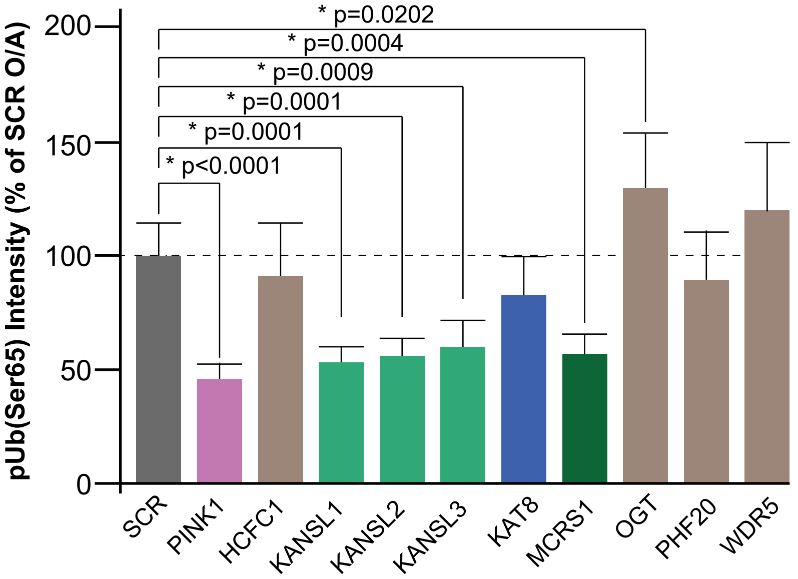


.

**Extended Data Figure 5 - Knockdown of the mitochondrial components of the NSL complex reduces pUb(Ser65) levels**
