## Extended Data Fig. 7 for "Regulation of mitophagy by the NSL complex underlies genetic risk for Parkinson’s disease at Chr16q11.2 and on the MAPT H1 allele"

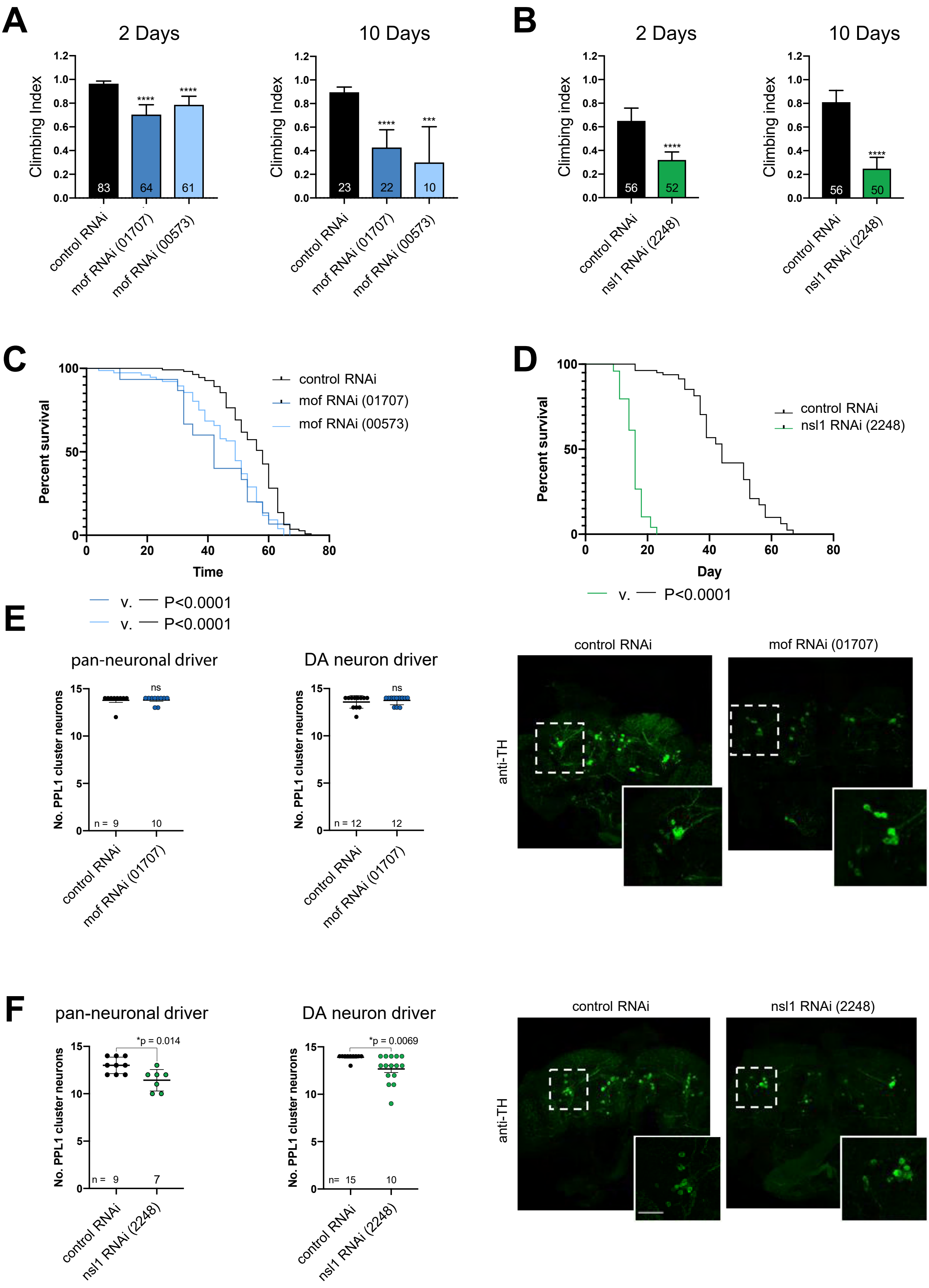


**Extended Data Figure 7 - Neuronal loss of *mof* or *nsl1* causes locomotor deficit, shortened lifespan and neurodegeneration**
