## Supplementary Table 1 for "Regulation of mitophagy by the NSL complex underlies genetic risk for Parkinson’s disease at Chr16q11.2 and on the MAPT H1 allele"

| **eQTL_dataset** | GTEx |
| --- | --- |
| **sequencing_method** | RNA-seq |
| **gene_symbol** | *KAT8* |
| **tissue** | Brain_Caudate_basal_ganglia |
| **braineac_probe_id** | NA |
| **nsnps** | 2499 |
| **PPH0** | 4.89E-10 |
| **PPH1** | 1.76E-07 |
| **PPH2** | 6.76E-04 |
| **PPH3** | 0.242293929 |
| **PPH4** | 0.75702972 |
| **PD_top_snp** | chr16:31000809 |
| **coloc_top_snp** | chr16:31048079 |
| **coloc_SNP_PPH4** | 0.08637809 |
| **coloc_eQTL_effect_allele** | T |
| **coloc_eQTL_other_allele** | C |
| **coloc_eQTL_beta** | -0.358743 |
| **coloc_eQTL_SE** | 0.0563187 |
| **coloc_eQTL_Freq1** | NA |
| **coloc_eQTL_p_val** | 3.49E-09 |
| **coloc_PD_Al1** | C |
| **coloc_PD_Al2** | T |
| **coloc_PD_beta** | -0.0739 |
| **coloc_PD_SE** | 0.0115 |
| **coloc_PD_Freq1** | 0.5972 |
| **coloc_PD_p_val** | 1.38E-10 |

**Supplementary Table 1. Results of the Colocalization analysis for *KAT8***.

PD_top_snp = lead SNP in the PD GWAS, coloc_top snp = Most likely SNP responsible for the colocalization signal, coloc_SNP_PPH4 = posterior probability of coloc top SNP being the true SNP responsible for the colocalization signal.
