## Supplementary Table 2 for "Regulation of mitophagy by the NSL complex underlies genetic risk for Parkinson’s disease at Chr16q11.2 and on the MAPT H1 allele"

| **ID** | *KAT8* | *KAT8* | *KAT8* | *KAT8* | *KAT8* | *KAT8* |
| --- | --- | --- | --- | --- | --- | --- |
| **CHR** | 16 | 16 | 16 | 16 | 16 | 16 |
| **P0** | 31127075 | 31127075 | 31127075 | 31127075 | 31128984 | 31127075 |
| **P1** | 31142714 | 31142714 | 31142714 | 31142714 | 31142714 | 31142714 |
| **HSQ** | 0.144 | 0.191 | 0.121 | 0.156 | 0.0641 | 0.207 |
| **BEST.GWAS.ID** | rs9938550 | rs9938550 | rs9938550 | rs9938550 | rs2305880 | rs9938550 |
| **BEST.GWAS.Z** | -6.82 | -6.82 | -6.82 | -6.82 | -6.84 | -6.82 |
| **EQTL.ID** | rs8046707 | rs1549293 | rs2855475 | rs12597511 | rs749767 | rs4527034 |
| **EQTL.R2** | 0.13718 | 0.129562 | 0.227501 | 0.19827 | 0.074695 | 0.00998 |
| **EQTL.Z** | -4.26 | -4.12 | -4.58 | -4.61 | -6.05 | -4.23 |
| **EQTL.GWAS.Z** | 6.28455 | 6.13675 | 6.1538 | 6.178 | 6.06897 | 6.1724 |
| **NSNP** | 240 | 240 | 240 | 240 | 248 | 240 |
| **NWGT** | 3 | 4 | 3 | 1 | 1 | 9 |
| **MODEL** | lasso | lasso | lasso | lasso | lasso | lasso |
| **MODELCV.R2** | 0.103 | 0.181407 | 0.17535 | 0.17222 | 0.07213 | 0.04964 |
| **MODELCV.PV** | 0.000889 | 1.37E-05 | 2.91E-05 | 2.56E-05 | 3.95E-09 | 0.0139 |
| **TWAS.Z** | -6.6932 | -6.2068 | -6.1876 | -6.178 | -6.069 | -6.04195 |
| **TWAS.P** | 2.18E-11 | 5.41E-10 | 6.11E-10 | 6.49E-10 | 1.29E-09 | 1.52E-09 |
| **FDR** | 2.09E-09 | 4.84E-08 | 7.15E-08 | 6.79E-08 | 4.37E-07 | 2.36E-07 |
| **Region** | GTEx.Brain _Cortex | GTEx.Brain_ Nucleus  accumbens _basal ganglia | GTEx.Brain_ Cerebellar Hemisphere | GTEx.Brain_ Frontal Cortex_BA9 | CMC.BRAIN .RNASEQ | GTEx.Brain_ Cerebellum |

**Supplementary Table 2. Results of the TWAS analysis for KAT8.**
