## Supplementary Table 3 for "Regulation of mitophagy by the NSL complex underlies genetic risk for Parkinson’s disease at Chr16q11.2 and on the MAPT H1 allele"

| **KAT name** | **Alternative name(s)** |
| --- | --- |
| KAT1 | HAT1 |
| KAT2A | GCN5 |
| KAT2B | PCAF |
| KAT3A | CREBBP, CBP |
| KAT3B | EP300 |
| KAT4 | TAF1, TFII250 |
| KAT5 | TIP60 |
| KAT6A | MOZ, MYST3 |
| KAT6B | MORF, MYST4 |
| KAT7 | HBO1, MYST2 |
| KAT8 | MOF, MYST1 |
| KAT9 | ELP3 |
| KAT12 | GTF3C4, TFIIIC90 |
| KAT13A | NCOA1, SRC1 |
| KAT13B | NCOA3, SRC3, ACTR |
| KAT13C | NCOA2 |
| KAT13D | CLOCK |
|  | ACAT1 |
|  | ATAT1 |
|  | ATF2 |
|  | BLOC1S1 (GCN5L1) |
|  | NAT10 |

**Supplementary Table 3. Complete list of the 22 KATs screened in the high content screen. (See Fig 1G)**
