## Supplementary Table 4 for "Regulation of mitophagy by the NSL complex underlies genetic risk for Parkinson’s disease at Chr16q11.2 and on the MAPT H1 allele"

| **Dunnett's multiple comparisons test** | **Mean Diff.** | **95.00% CI of diff.** | **Significant?** | **Summary** | **Adjusted P Value** |
| --- | --- | --- | --- | --- | --- |
| DMSO |  |  |  |  |  |
| SCR vs. KAT8 | 0 | -14.19 to 14.19 | No | ns | >0.9999 |
| SCR vs. KANSL1 | 0 | -14.19 to 14.19 | No | ns | >0.9999 |
| SCR vs. PINK1 | 0 | -14.19 to 14.19 | No | ns | >0.9999 |
| 1 h |  |  |  |  |  |
| SCR vs. KAT8 | 21 | 6.807 to 35.19 | Yes | ** | 0.0017 |
| SCR vs. KANSL1 | 30 | 15.81 to 44.19 | Yes | **** | <0.0001 |
| SCR vs. PINK1 | 38 | 23.81 to 52.19 | Yes | **** | <0.0001 |
| 2 h |  |  |  |  |  |
| SCR vs. KAT8 | 30 | 15.81 to 44.19 | Yes | **** | <0.0001 |
| SCR vs. KANSL1 | 53 | 38.81 to 67.19 | Yes | **** | <0.0001 |
| SCR vs. PINK1 | 71 | 56.81 to 85.19 | Yes | **** | <0.0001 |
| 3 h |  |  |  |  |  |
| SCR vs. KAT8 | 33 | 18.81 to 47.19 | Yes | **** | <0.0001 |
| SCR vs. KANSL1 | 51 | 36.81 to 65.19 | Yes | **** | <0.0001 |
| SCR vs. PINK1 | 73 | 58.81 to 87.19 | Yes | **** | <0.0001 |
| 4 h |  |  |  |  |  |
| SCR vs. KAT8 | 15 | 0.8065 to 29.19 | Yes | * | 0.0355 |
| SCR vs. KANSL1 | 28 | 13.81 to 42.19 | Yes | **** | <0.0001 |
| SCR vs. PINK1 | 49 | 34.81 to 63.19 | Yes | **** | <0.0001 |
| 5 h |  |  |  |  |  |
| SCR vs. KAT8 | 9 | -5.193 to 23.19 | No | ns | 0.307 |
| SCR vs. KANSL1 | 21 | 6.807 to 35.19 | Yes | ** | 0.0017 |
| SCR vs. PINK1 | 39 | 24.81 to 53.19 | Yes | **** | <0.0001 |
| 6 h |  |  |  |  |  |
| SCR vs. KAT8 | 14 | -0.1935 to 28.19 | No | ns | 0.0542 |
| SCR vs. KANSL1 | 24 | 9.807 to 38.19 | Yes | *** | 0.0003 |
| SCR vs. PINK1 | 42 | 27.81 to 56.19 | Yes | **** | <0.0001 |
| 7 h |  |  |  |  |  |
| SCR vs. KAT8 | 19 | 4.807 to 33.19 | Yes | ** | 0.0051 |
| SCR vs. KANSL1 | 23 | 8.807 to 37.19 | Yes | *** | 0.0005 |
| SCR vs. PINK1 | 40 | 25.81 to 54.19 | Yes | **** | <0.0001 |

| Within each row, compare columns (simple effects within rows) | |
| --- | --- |
| Number of families | 8 |
| Number of comparisons | 3 |
| Alpha | 0.5 |

**Supplementary Table 4. p-values for Figure 3B.**
