## Supplementary Table 5 for "Regulation of mitophagy by the NSL complex underlies genetic risk for Parkinson’s disease at Chr16q11.2 and on the MAPT H1 allele"

| Within each row, compare columns (simple effects within rows) | | | |  |  |
| --- | --- | --- | --- | --- | --- |
| Number of families | 5 |  |  |  |  |
| Number of comparisons per family | 3 |  |  |  |  |
| Alpha | 0.05 |  |  |  |  |
| **Dunnett's multiple**  **comparisons test** | **Mean**  **Diff.** | **95.00% CI of diff.** | **Below**  **threshold?** | **Summary** | **Adjusted**  **P Value** |
| 0 h |  |  |  |  |  |
| SCR OA vs. KAT8 OA | -0.003343 | -0.04335 to 0.03667 | No | ns | 0.9940 |
| SCR OA vs. KANSL1 OA | -0.002681 | -0.04269 to 0.03733 | No | ns | 0.9969 |
| SCR OA vs. PINK1 OA | 0.004262 | -0.03575 to 0.04427 | No | ns | 0.9878 |
| 2 h |  |  |  |  |  |
| SCR OA vs. KAT8 OA | 0.04496 | 0.004950 to 0.08497 | Yes | * | 0.0243 |
| SCR OA vs. KANSL1 OA | 0.04991 | 0.009896 to 0.08992 | Yes | * | 0.0113 |
| SCR OA vs. PINK1 OA | 0.05799 | 0.01798 to 0.09800 | Yes | ** | 0.0029 |
| 4 h |  |  |  |  |  |
| SCR OA vs. KAT8 OA | 0.06233 | 0.02232 to 0.1023 | Yes | ** | 0.0014 |
| SCR OA vs. KANSL1 OA | 0.07349 | 0.03348 to 0.1135 | Yes | *** | 0.0002 |
| SCR OA vs. PINK1 OA | 0.07958 | 0.03957 to 0.1196 | Yes | **** | <0.0001 |
| 6 h |  |  |  |  |  |
| SCR OA vs. KAT8 OA | 0.05593 | 0.01592 to 0.09594 | Yes | ** | 0.0042 |
| SCR OA vs. KANSL1 OA | 0.06001 | 0.02000 to 0.1000 | Yes | ** | 0.0021 |
| SCR OA vs. PINK1 OA | 0.05635 | 0.01634 to 0.09636 | Yes | ** | 0.0039 |
| 8 h |  |  |  |  |  |
| SCR OA vs. KAT8 OA | 0.04123 | 0.001219 to 0.08124 | Yes | * | 0.0421 |
| SCR OA vs. KANSL1 OA | 0.03317 | -0.006843 to 0.07318 | No | ns | 0.1231 |
| SCR OA vs. PINK1 OA | 0.02560 | -0.01441 to 0.06561 | No | ns | 0.2876 |

**Supplementary Table 5. p-values for Figure 4B.**
