## Supplementary Table 6 for "Regulation of mitophagy by the NSL complex underlies genetic risk for Parkinson’s disease at Chr16q11.2 and on the MAPT H1 allele"

| **hetSNP** | **Sample ID** | **Read count** | **Allele** | **Ens.Alt** | **Individual ID** | **Tissue** |
| --- | --- | --- | --- | --- | --- | --- |
| 17:44108355 | A653_043 | 47 | A | A | 004_06 | SNIG |
| 17:44108355 | A653_043 | 20 | G | A | 004_06 | SNIG |
| 17:44108355 | A653_441 | 69 | A | A | 032_09 | PUTM |
| 17:44108355 | A653_441 | 0 | G | A | 032_09 | PUTM |
| 17:44159849 | A653_031 | 2 | C | C | 024_09 | PUTM |
| 17:44159849 | A653_031 | 23 | T | C | 024_09 | PUTM |
| 17:44159849 | A653_719 | 58 | C | C | 035_09 | SNIG |
| 17:44159849 | A653_719 | 26 | T | C | 035_09 | SNIG |
| 17:44248769 | A653_093 | 36 | C | C | 004_08 | SNIG |
| 17:44248769 | A653_093 | 4 | T | C | 004_08 | SNIG |
| 17:44248769 | A653_184 | 30 | C | C | 013_09 | PUTM |
| 17:44248769 | A653_184 | 7 | T | C | 013_09 | PUTM |
| 17:44248769 | A653_326 | 26 | C | C | 017_09 | PUTM |
| 17:44248769 | A653_326 | 3 | T | C | 017_09 | PUTM |
| 17:44248769 | A653_617 | 22 | C | C | 029_09 | SNIG |
| 17:44248769 | A653_617 | 0 | T | C | 029_09 | SNIG |
| 17:44248769 | A653_679 | 23 | C | C | 030_06 | PUTM |
| 17:44248769 | A653_679 | 3 | T | C | 030_06 | PUTM |
| 17:44248769 | A653_753 | 43 | C | C | 029_09 | PUTM |
| 17:44248769 | A653_753 | 10 | T | C | 029_09 | PUTM |
| 17:44248769 | A653_794 | 24 | C | C | 036_09 | SNIG |
| 17:44248769 | A653_794 | 4 | T | C | 036_09 | SNIG |
| 17:44248814 | A653_043 | 10 | A | A | 004_06 | SNIG |
| 17:44248814 | A653_043 | 0 | G | A | 004_06 | SNIG |
| 17:44248814 | A653_326 | 19 | A | A | 017_09 | PUTM |
| 17:44248814 | A653_326 | 3 | G | A | 017_09 | PUTM |
| 17:44248814 | A653_617 | 12 | A | A | 029_09 | SNIG |
| 17:44248814 | A653_617 | 0 | G | A | 029_09 | SNIG |
| 17:44248814 | A653_679 | 19 | A | A | 030_06 | PUTM |
| 17:44248814 | A653_679 | 2 | G | A | 030_06 | PUTM |
| 17:44248814 | A653_753 | 25 | A | A | 029_09 | PUTM |
| 17:44248814 | A653_753 | 5 | G | A | 029_09 | PUTM |
| 17:44248814 | A653_950 | 28 | A | A | 006_10 | SNIG |
| 17:44248814 | A653_950 | 8 | G | A | 006_10 | SNIG |

| **Supplementary Table 6. Summary of the results of ASE analysis across the *KANSL1* gene.** | | |
| --- | --- | --- |
| **Column name** | | **Description** |
| hetSNP |  | Heterozygous SNP with average read depth >15 across samples |
| Avg.Reads.all.samples | | Average read depth across all samples |
| min.FDR |  | Minimum false discovery rate across the samples |
| Allele1 |  | Allele 1 for the hetSNP |
| Allele2 |  | Allele 2 for the hetSNP |
| ens.alt |  | Ensembl alternate allele |
| symbol |  | Gene name obtained using the variant effect predictor tool (VEP) |
| most.severe.consequence | | Most severe of all observed consequence types reported for the hetSNP (VEP) |
| ASE |  | Indicates ASE = 'Y' if min.FDR < 0.05, 'N' if min.FDR >=0.05 |
| **The following column details were from obtained from LDProxy (https://ldlink.nci.nih.gov/)** | | |
| RSID |  | Identifier for the hetSNP |
| Alleles |  | SNP alleles |
| MAF |  | Minor allele frequency |
| Dprime |  | Indicator of allelic segregation for two genetic variants |
| R2 |  | Measure of correlation of alleles for two genetic variants. |
