## Supplementary Table 7 for "Regulation of mitophagy by the NSL complex underlies genetic risk for Parkinson’s disease at Chr16q11.2 and on the MAPT H1 allele"

| **hetSNP** | **Sample ID** | **Read count** | **Allele** | **Ens.Alt** | **Individual ID** | **Tissue** |
| --- | --- | --- | --- | --- | --- | --- |
| 17:44067400 | A653_031 | 18 | C | C | 024_09 | PUTM |
| 17:44067400 | A653_031 | 3 | T | C | 024_09 | PUTM |
| 17:44102689 | A653_031 | 5 | C | C | 024_09 | PUTM |
| 17:44102689 | A653_031 | 21 | G | C | 024_09 | PUTM |
| 17:44103616 | A653_031 | 23 | C | T | 024_09 | PUTM |
| 17:44103616 | A653_031 | 0 | T | T | 024_09 | PUTM |
| 17:44103825 | A653_031 | 11 | C | C | 024_09 | PUTM |
| 17:44103825 | A653_031 | 31 | T | C | 024_09 | PUTM |
| 17:44104509 | A653_043 | 43 | C | C | 004_06 | SNIG |
| 17:44104509 | A653_043 | 82 | T | C | 004_06 | SNIG |
| 17:44102604 | A653_056 | 85 | C | C | 015_07 | PUTM |
| 17:44102604 | A653_056 | 138 | T | C | 015_07 | PUTM |
| 17:44104343 | A653_093 | 84 | A | C | 004_08 | SNIG |
| 17:44104343 | A653_093 | 34 | C | C | 004_08 | SNIG |
| 17:44102689 | A653_171 | 37 | C | C | 040_08 | SNIG |
| 17:44102689 | A653_171 | 87 | G | C | 040_08 | SNIG |
| 17:44039691 | A653_177 | 118 | A | G | 015_07 | SNIG |
| 17:44039691 | A653_177 | 184 | G | G | 015_07 | SNIG |
| 17:44104343 | A653_177 | 35 | C | C | 015_07 | SNIG |
| 17:44104343 | A653_177 | 74 | A | C | 015_07 | SNIG |
| 17:44102689 | A653_205 | 30 | C | C | 040_08 | PUTM |
| 17:44102689 | A653_205 | 64 | G | C | 040_08 | PUTM |
| 17:44102689 | A653_225 | 36 | C | C | 004_08 | PUTM |
| 17:44102689 | A653_225 | 70 | G | C | 004_08 | PUTM |
| 17:44104343 | A653_243 | 37 | C | C | 021_09 | SNIG |
| 17:44104343 | A653_243 | 71 | A | C | 021_09 | SNIG |
| 17:44102689 | A653_283 | 47 | C | C | 017_09 | SNIG |
| 17:44102689 | A653_283 | 90 | G | C | 017_09 | SNIG |
| 17:44102865 | A653_283 | 79 | A | C | 017_09 | SNIG |
| 17:44102865 | A653_283 | 41 | C | C | 017_09 | SNIG |
| 17:44104343 | A653_283 | 42 | C | C | 017_09 | SNIG |
| 17:44104343 | A653_283 | 84 | A | C | 017_09 | SNIG |
| 17:44104509 | A653_283 | 85 | T | C | 017_09 | SNIG |
| 17:44104509 | A653_283 | 43 | C | C | 017_09 | SNIG |
| 17:44067400 | A653_288 | 50 | C | C | 038_08 | PUTM |
| 17:44067400 | A653_288 | 12 | T | C | 038_08 | PUTM |
| 17:44104509 | A653_326 | 73 | T | C | 017_09 | PUTM |
| 17:44104509 | A653_326 | 33 | C | C | 017_09 | PUTM |
| 17:44101563 | A653_441 | 34 | C | C | 032_09 | PUTM |
| 17:44101563 | A653_441 | 0 | T | C | 032_09 | PUTM |
| 17:44102638 | A653_441 | 0 | A | G | 032_09 | PUTM |
| 17:44102638 | A653_441 | 197 | G | G | 032_09 | PUTM |
| 17:44102689 | A653_441 | 77 | C | C | 032_09 | PUTM |
| 17:44102689 | A653_441 | 0 | G | C | 032_09 | PUTM |
| 17:44103296 | A653_441 | 112 | C | C | 032_09 | PUTM |
| 17:44103296 | A653_441 | 0 | T | C | 032_09 | PUTM |
| 17:44103825 | A653_441 | 75 | C | C | 032_09 | PUTM |
| 17:44103825 | A653_441 | 0 | T | C | 032_09 | PUTM |
| 17:44103826 | A653_441 | 74 | A | A | 032_09 | PUTM |
| 17:44103826 | A653_441 | 0 | G | A | 032_09 | PUTM |
| 17:44067400 | A653_627 | 9 | T | C | 033_09 | PUTM |
| 17:44067400 | A653_627 | 30 | C | C | 033_09 | PUTM |
| 17:44068924 | A653_627 | 39 | A | A | 033_09 | PUTM |
| 17:44068924 | A653_627 | 72 | G | A | 033_09 | PUTM |
| 17:44102933 | A653_627 | 73 | C | C | 033_09 | PUTM |
| 17:44102933 | A653_627 | 118 | T | C | 033_09 | PUTM |
| 17:44104343 | A653_627 | 68 | A | C | 033_09 | PUTM |
| 17:44104343 | A653_627 | 35 | C | C | 033_09 | PUTM |
| 17:44104509 | A653_627 | 92 | T | C | 033_09 | PUTM |
| 17:44104509 | A653_627 | 53 | C | C | 033_09 | PUTM |
| 17:44102689 | A653_643 | 24 | C | C | 017_08 | PUTM |
| 17:44102689 | A653_643 | 54 | G | C | 017_08 | PUTM |
| 17:44104509 | A653_679 | 11 | C | C | 030_06 | PUTM |
| 17:44104509 | A653_679 | 40 | T | C | 030_06 | PUTM |
| 17:44067400 | A653_719 | 13 | T | C | 035_09 | SNIG |
| 17:44067400 | A653_719 | 51 | C | C | 035_09 | SNIG |
| 17:44102689 | A653_719 | 45 | C | C | 035_09 | SNIG |
| 17:44102689 | A653_719 | 84 | G | C | 035_09 | SNIG |
| 17:44067400 | A653_738 | 23 | C | C | 036_09 | PUTM |
| 17:44067400 | A653_738 | 3 | T | C | 036_09 | PUTM |
| 17:44067400 | A653_753 | 24 | C | C | 029_09 | PUTM |
| 17:44067400 | A653_753 | 6 | T | C | 029_09 | PUTM |
| 17:44101563 | A653_763 | 0 | C | C | 034_08 | PUTM |
| 17:44101563 | A653_763 | 33 | T | C | 034_08 | PUTM |
| 17:44103296 | A653_763 | 0 | C | C | 034_08 | PUTM |
| 17:44103296 | A653_763 | 91 | T | C | 034_08 | PUTM |
| 17:44068924 | A653_794 | 39 | A | A | 036_09 | SNIG |
| 17:44068924 | A653_794 | 73 | G | A | 036_09 | SNIG |

| **Supplementary Table 7. Summary of the results of ASE analysis across the *MAPT* gene.** | | |
| --- | --- | --- |
| **Column name** |  | **Description** |
| hetSNP |  | Heterozygous SNP with average read depth >15 across samples |
| Avg.Reads.all.samples | | Average read depth across all samples |
| min.FDR |  | Minimum false discovery rate across the samples |
| Allele1 |  | Allele 1 for the hetSNP |
| Allele2 |  | Allele 2 for the hetSNP |
| ens.alt |  | Ensembl alternate allele |
| symbol |  | Gene name obtained using the variant effect predictor tool (VEP) |
| most.severe.consequence | | Most severe of all observed consequence types reported for the hetSNP (VEP) |
| ASE |  | Indicates ASE = 'Y' if min.FDR < 0.05, 'N' if min.FDR >=0.05 |
| **The following column details were from obtained from LDProxy (https://ldlink.nci.nih.gov/)** | | |
| RSID |  | Identifier for the hetSNP |
| Alleles |  | SNP alleles |
| MAF |  | Minor allele frequency |
| Dprime |  | Indicator of allelic segregation for two genetic variants |
| R2 |  | Measure of correlation of alleles for two genetic variants. |
