## Supplementary Table 8 for "Regulation of mitophagy by the NSL complex underlies genetic risk for Parkinson’s disease at Chr16q11.2 and on the MAPT H1 allele"

| **17q21 ORFs** | |
| --- | --- |
| *ACBD4* | *HEXIM1* |
| *ADAM11* | *HIGD1B* |
| *ARHGAP27* | *KANSL1* |
| *ARL17A* | *KIF18B* |
| *ARL17B* | *LRRC37A* |
| *C1QL1* | *LRRC37A2* |
| *CCDC103* | *MAP3K14* |
| *CDC27* | *MAPT* |
| *CRHR1* | *MYL4* |
| *DBF4B* | *NMT1* |
| *DCAKD* | *NSF* |
| *EFTUD2* | *PLEKHM1* |
| *FMNL1* | *RPRML* |
| *GFAP* | *SPPL2C* |
| *GJC1* | *STH* |
| *GOSR2* | *WNT3* |

**Supplementary Table 8. Complete list of the 32 ORFs in the 17q21 locus screened in the high content screen (See Fig 5).**
