## Supplementary Table 9 for "Regulation of mitophagy by the NSL complex underlies genetic risk for Parkinson’s disease at Chr16q11.2 and on the MAPT H1 allele"

|  | **Sequence 5'-3'** | |  |
| --- | --- | --- | --- |
| **Gene Target** | **Forward** | **Reverse** | **Product Size / bp** |
| *RPL18A* | CCCACAACATGTACCGGGAA | TCTTGGAGTCGTGGAACTGC | 180 |
| *KANSL1* | ATCCTCCACACAGTCCCTTG | CCCCTTCTCCTCCTTACTGG | 121 |
| *KAT8* | TCACTCGCAACCAAAAGCG | GATCGCCTCATGCTCCTTCT | 107 |
| *PINK1* | GTGGAACATCTCGGCAGGTT | CCTCTCTTGGATTTTCTGTAAGTGAC | 129 |

**Supplementary Table 9. List of primer pairs used for RT-qPCR of target genes.**
